# EXPRESSION OF CONJUGATION GENES IS CONTROLLED BY PROCESSIVE ANTITERMINATION AND A NOVEL ZIPPER-TYPE TRANSCRIPTIONAL ATTENUATION MECHANISM

**DOI:** 10.1101/2025.11.24.689506

**Authors:** Daniel González Álvarez, Andrés Miguel-Arribas, Sandeepani Ranaweera, Fernando Freire-Gómez, David Abia, Anne de Jong, Ling Juan Wu, Paul Babitzke, Wilfried J.J. Meijer

## Abstract

Proper expression of genes clustered in operons, particularly large operons, can be complex, often consisting of multiple regulatory switches. The conjugation operon present on the *Bacillus subtilis* conjugative plasmid pLS20 is over 32 kb long. This operon starts with a 456 nt leader region, which is followed by the first two genes of the operon, encoding a two-component processive antitermination system. Here, we demonstrate that the long leader region encodes a transcriptional attenuator that we named cATT_pLS20_. In vivo and in vitro analyses showed that the attenuator is composed of three segments: a stem-loop structure with a long imperfect stem that is preceded by a sequence that may form a weak stem-loop and followed by a sequence that may form an intrinsic terminator. Sequences of the upstream stem loop and downstream terminator are complementary, allowing further extension of the long stem, thereby generating an antiterminator conformation. Based on similarity with a zipper we coined this the zipper-type attenuator. Similar zipper-type attenuators are present upstream of the first genes of conjugation operons of all pLS20 family plasmids, as well as on many other conjugative plasmids in Gram-positive bacteria, suggesting that a common attenuation mechanism regulates expression of many conjugative operons.

**GRAPHICAL ABSTRACT:** 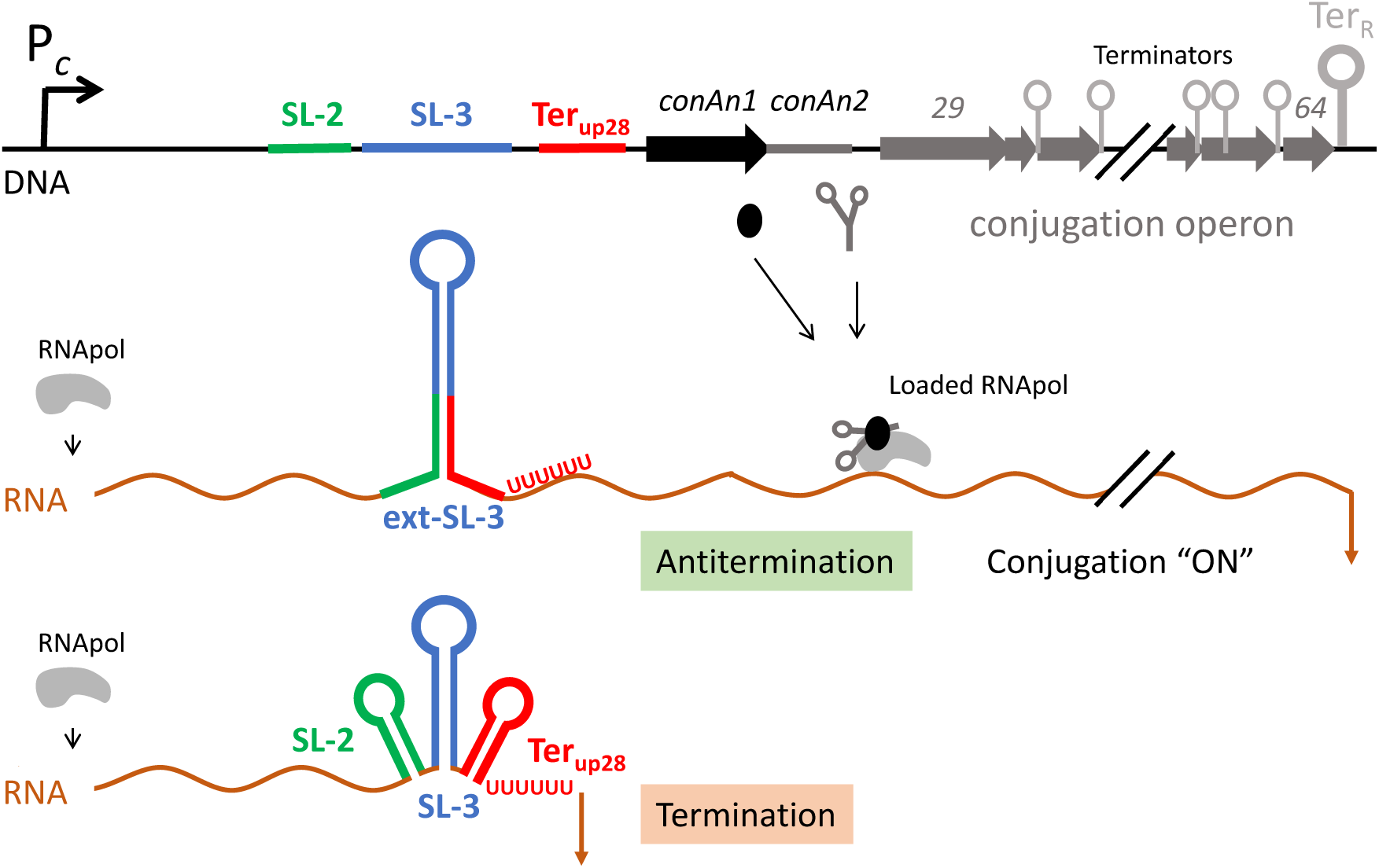

## INTRODUCTION

Gene expression is a highly dynamic process with only a portion of the genome expressed at any given time. The expression profile can change rapidly and profoundly in response to changing environmental conditions. Multiple mechanisms acting on the three stages of the transcription cycle - initiation, elongation, and termination - have evolved to achieve these adaptive gene expression profiles (1–11).

Bacterial conjugation is the process by which a genetic element, named the conjugative element, can transfer from a donor to a recipient cell via a channel that connects both cells (12–14). Often a conjugative element is located on a plasmid, which is then called a conjugative plasmid. In addition to the metabolic burden, expression of conjugation genes significantly affects the structure and the physiology of the bacterial host. For instance, some of the conjugation genes encode cell wall associated adhesins and integral membrane proteins that change characteristics of the bacterial cell wall and membrane. The adhesins facilitate attachment of donor to recipient cells, while the membrane proteins organise into a membrane-embedded type IV secretions system (T4SS) that is responsible for transferring a ssDNA strand of the plasmid into the recipient cell (14–17). The conjugation process also affects the biology of the plasmid such as the mode of DNA replication. At the onset of conjugation, the plasmid switches from the common theta replication mechanism used to duplicate the plasmid to a rolling-circle mechanism to generate the ssDNA strand that is transferred to the recipient cell. It is therefore not surprising that expression of conjugation genes are tightly controlled (18–20), which is the case for the *Bacillus subtilis* conjugative plasmid pLS20 (21). This 64 kb plasmid contains a large operon of >32 kb that comprises probably all the conjugation genes, which are numbered from gene *28* to *64*. We previously identified regulatory mechanisms that control (i) the initiation of transcription at the single strong conjugation promoter P*_c_*, located upstream of the first gene of the conjugation operon (gene *28a*), and (ii) processive transcription of the extremely long conjugation operon. Initiation of transcription at the conjugation promoter P*_c_* involves multiple layers of control, involving three proteins encoded by genes *25*-*27* located immediately upstream of the conjugation operon. Gene *27*, whose orientation is divergent to the genes in the conjugation operon, encodes a transcriptional repressor named Rco, while genes *25* and *26* form a bicistronic operon encoding Rap, the Rco antirepressor, and a small prepropeptide Phr, respectively. Rco tetramers can bind two operators, one of which overlaps the divergently oriented promoters P*_c_* and P*_r_*, the latter driving expression of Rco (Figure 1A). When bound to its operators, Rco activates expression of the weak P*_r_* promoter and tightly represses the strong P*_c_* promoter, keeping conjugation in the default “off” state. Transcription from P*_c_*is induced when Rap binds to and inactivates Rco. After secretion into the medium and a second proteolytic step, the pentapeptide Phr* corresponding to the five C-terminal residues of Phr is generated, which acts as a signal peptide. When imported by donor cells, Phr* interacts with Rap thereby inactivating its antirepressor function, and allowing Rco to reestablish repression of the conjugation operon. High Phr* levels are generated only when all or most cells in a population contain pLS20. Therefore, this quorum sensing mechanism allows expression of the conjugation genes only when donor cells are potentially surrounded by plasmid-free recipient cells. The three proteins Rap, Phr* and Rco have been characterized at considerable detail at the functional, biochemical, biophysical, and structural levels (22–26).

**Figure 1.**
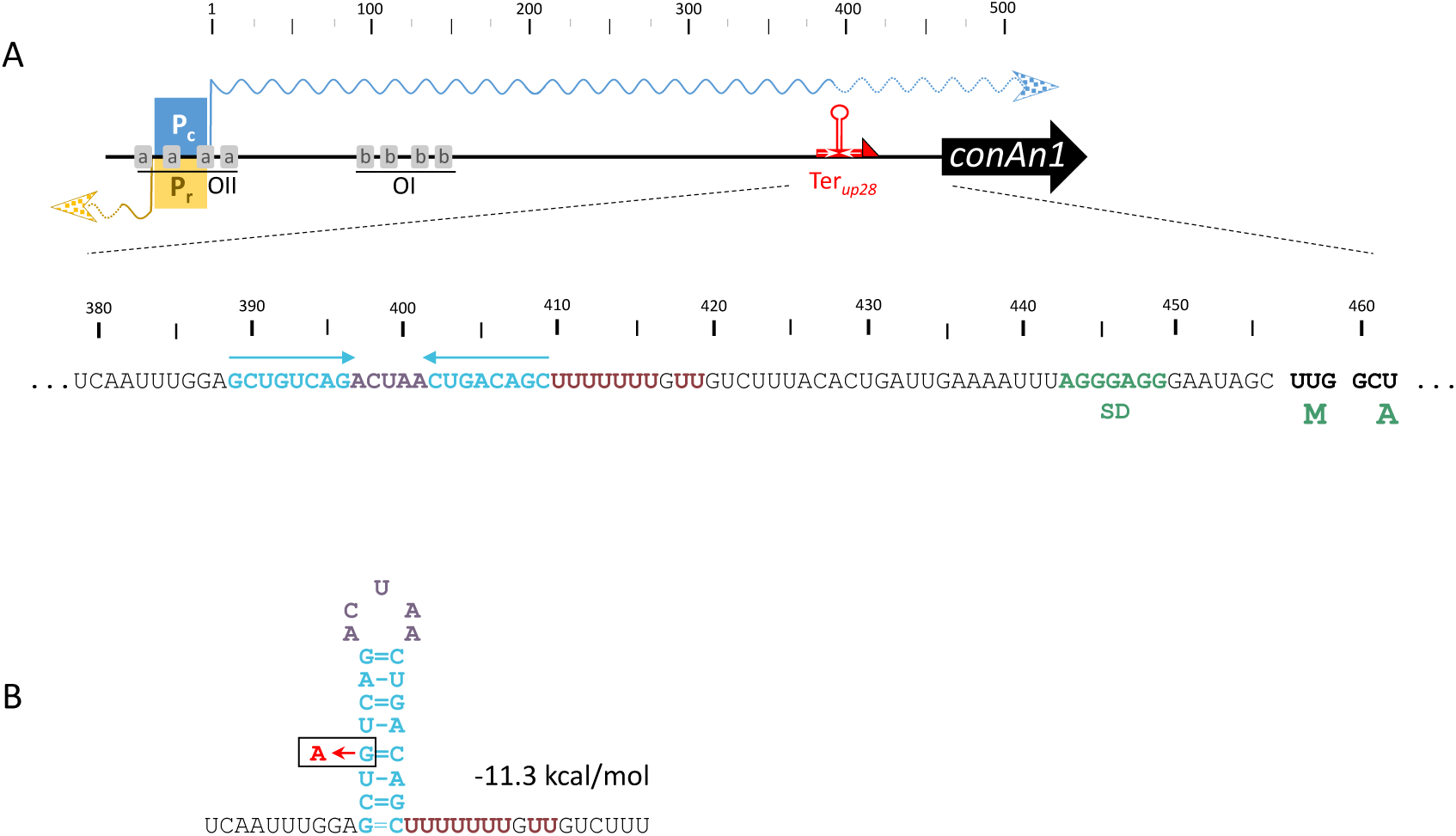
An intrinsic terminator (Ter*_up28_*) precedes *conAn1,* the first gene of the conjugation operon. **(A)** Schematic diagram of the intergenic region between the conjugation promoter P*_c_*, and the first gene of the conjugation operon, *conAn1* (black arrow). The positions of the divergent promoters P*_r_* and P*_c_* are indicated with a yellow and blue box, respectively. Transcription start sites are indicated with vertical lines. Transcripts derived from P*_r_* and P*_c_* are indicated with yellow and blue wavy lines, respectively. Numbering shown at the top corresponds to the P*_c_*-derived transcript; position 1 is the transcription start site. Rco binding sites in the Rco operators OII and OI, are indicated with grey boxes labelled a and b, respectively (23). The putative intrinsic terminator Ter*_up28_* is indicated. The sequence of the region encompassing Ter*_up28_* is shown. Sequences forming the predicted stem, loop, and U-rich tract of the terminator are shown in blue, purple and red, respectively. Sequences that would form the stem are also marked with blue arrows. The likely Shine-Dalgarno (SD) sequence and the first two codons of *conAn1* are indicated in green. **(B)** Predicted structure of Ter*_up28_*. The predicted thermodynamic stability is indicated. The colour code to indicate sequences forming the stem, loop, and U-rich tract is the same as those used in panel A. The position of the G to A point mutation in the 5′ stem of Ter*_up28_*present in strain DGA06 is indicated.

The first two genes of the conjugation operon, genes *28a* and *28b*, code for a two-component processive antitermination (P-AT) system, named *conAn* (27). Gene *28a* encodes protein ConAn1 and gene *28b* encodes an RNA, named ConAn2. Whereas ConAn2 exerts antitermination, ConAn1 acts as a processivity factor. The *conAn* antitermination system functions only in *cis*. The association of ConAn1 and ConAn2 with RNA polymerase (RNAP) that has initiated transcription at P*_c_* alters the elongation complex such that it is capable of bypassing at least six, and probably up to 20 intrinsic terminators, that are distributed along the conjugation operon. A specialized intrinsic terminator with an exceptionally long stem that is resistant to *conAn*-mediated antitermination demarcates the end of the conjugation operon (28).

Here, we show that in addition to these mechanisms, the conjugation genes of pLS20 are controlled by a transcriptional attenuator that is located in the 456 nt leader region upstream of *conAn1*. Thus, conjugation gene transcription is controlled by attenuation and P-AT mechanisms that act at a single and at multiple intrinsic terminators, respectively. Importantly, we found that a transcriptional attenuator similar to that on pLS20 precedes the conjugation operons of other pLS20 family plasmids, and many other plasmids replicating in Gram+ bacteria, suggesting that conjugation is commonly controlled by these dual post-transcription initiation control mechanisms.

## MATERIAL AND METHODS

### Bacterial strains, plasmids, media and oligonucleotides

*Escherichia coli* and *B. subtilis* strains were cultured in Lysogeny Broth (LB) medium (29). Cultures were grown either in liquid medium with shaking or on 1.5% LB agar plates at 37 °C. When required, antibiotics were added as follows: ampicillin (100 µg/mL) for *E. coli*; spectinomycin (100 µg/mL) and chloramphenicol (5 µg/mL) for *B. subtilis*. All *B. subtilis* strains used in this study were isogenic with strain 168. A list of bacterial strains is provided in Table S1. Plasmids and oligonucleotides are listed in Tables S2 and S3, respectively. All oligonucleotides were synthesized by Integrated DNA Technologies (IDT).

### Transformation

*E. coli* competent cells were prepared and transformed using standard methods (30). Generation of competent *B. subtilis* cells and transformations were prepared as described previously (31).

### Construction of plasmids and strains

DNA techniques were performed using standard molecular methods (30). Plasmids were isolated from *E. coli* strains using the NZYMiniprep kit (NZYtech). PCR fragments were purified using NZYGelpure (NZYtech). In some cases, complementary primers were hybridized, and the resulting product was used as an insert in ligation reactions. In these cases, a 200 μL mixture was prepared (50 mM NaCl and 40 mM Tris-HCl, pH 7.5) containing 2,000 pmol of each primer. After boiling, the solution was slowly cooled to room temperature. The hybridized oligonucleotides were then used as an insert in ligation reactions with plasmid pAND101 (27) digested with SalI and NheI. All enzymes used were purchased from New England Biolabs, USA (NEB). Cloned fragments were confirmed by DNA sequencing. The Q5 site-directed mutagenesis kit (NEB) was used for introducing point mutations in the attenuator region of strains DGA73, DGA84, and DGA86.

Specific information of the construction of each plasmid, including the oligonucleotides and restriction enzymes used to amplify and clone the fragments, is presented in supplementary Table S2. Oligonucleotides used are listed in supplementary Table S3. Derivatives of vector pAND101 were used to test terminator or promoter activity in vivo. First, a PCR fragment containing a putative terminator was cloned at the multiple cloning site (MCS) located between the P*_spank_* promoter and *gfp* on plasmid pAND101. The resulting plasmid was linearized and the P*_spank_*-[insert]-*gfp* construct was integrated into the *amyE* locus by transformation. The resulting strains were used to study terminator or promoter activity of the cloned region by growing cells in the presence or absence of IPTG, respectively.

Marker less in-frame deletions on pLS20cat were constructed making use of the pMiniMAD2 vector (32). The method is based on single cross-over integration of a temperature–sensitive replicon via homologous recombination at the restrictive temperature, followed by permitting replication of the integrated replicon by growth at the permissive temperature to facilitate deletion of the replicon, thereby generating the desired deletion (33). Briefly, ⁓600 bp of DNA upstream and downstream of the region of interest were amplified by PCR, which were then joined in a second step by overlapping PCR. The resulting fused PCR product was purified, digested with NdeI and BamHI, and then cloned into the pMiniMAD2 vector digested with the same enzymes. For a full description of the strategy see (31). More details regarding the construction of plasmids are described in Supplemental Tables S2 and S3.

### Flow cytometry

Bacteria cultures were grown overnight (ON) at 37 °C with shaking in LB medium and diluted 1:100 into fresh LB the following day. Cultures were incubated at 37 °C with shaking until reaching an OD₆₀₀ of 0.8–1.0. Cells were harvested by centrifugation in 2 mL microcentrifuge tubes and washed twice with phosphate-buffered saline (PBS; 137 mM NaCl, 2.7 mM KCl, 10 mM Na₂HPO₄, 1.8 mM KH₂PO₄). The final pellet was resuspended in 2 mL PBS.

Fluorescence was measured using a FACSCalibur flow cytometer (Becton Dickinson, USA) equipped with a 488-nm argon laser. For each sample, at least 1×10⁵ events were collected using a 530/30-nm band-pass filter. Data acquisition was performed with CellQuest Pro (Becton Dickinson), and analyses were carried out using FlowJo v6.4.1 (TreeStar, USA). Fluorescence values are reported as the mean of the geometric mean (Geomean) fluorescence intensities obtained from ≥100,000 cells per sample across three independent biological replicates.

### Bioinformatics analysis of sequences upstream of conjugation operons

The 35 pLS20 family of plasmids (34) together with newly identified plasmids pBcaBIK3 and pBsuNRS6190 (35) were screened for the presence of a putative attenuator upstream of their conjugation operon. For this analysis, the 280 bp region upstream of the *conAn1* homolog were extracted and identical sequences were purged resulting in 19 unique sequences. These sequences were used to (i) generate a multiple sequence alignment with LocARNA using standard settings (36), and (ii) determine the phylogenetic relationship by constructing a maximum-likelihood tree with IQ-TREE software (37) version 2.2.6 using the edge-linked model (38) and applying the following settings: automatic selection of model, 1000 bootstrapping and 1000 replicates to perform SH-like approximate likelihood ratio test.

The following strategy was used to identify putative attenuators on conjugative non-pLS20 family plasmids of Gram-positive bacteria. Plasmid sequences deposited in the plasmid database (39,40) were screened for the presence of a *conAn1* homolog that was followed downstream by typical conjugation genes like ATPase encoding genes *virB4* and *virD4*, resulting in 1,162 hits. The 500 bp region immediately upstream of the *conAn1* gene of each plasmid was extracted. After purging duplicates and removing sequences belonging to plasmids of the pLS20 family, 423 unique sequences were obtained. According to the ARNold webserver (41,42), 339 of these sequences were predicted to contain a putative terminator (see supplemental Table S4). Strikingly, in many cases the predicted terminator was located closely upstream of the *conAn1* gene, as in the case of the pLS20 family plasmids. The presence of a terminator upstream of the first gene of a long conjugation operon suggests it might function as an attenuator.

### In vitro termination assays

In vitro transcription was performed by modifying published procedures (43–45). A double-stranded gBlock gene fragment was designed as the template for multi-round *in vitro* transcription experiments. This fragment consisted of a strong engineered promoter and the pLS20 conjugative operon leader region from positions 199-466 with respect to the transcription start site (TSS). Primers were designed to PCR amplify the gBlock fragment to use as the template for in vitro transcription. The RNAP open complex was assembled by mixing equal volumes of 2X template (100 nM) and 2X RNAP master mix (100 μg/ml bovine serum albumin, 150 μg/ml *B. subtilis* RNAP holoenzyme, and 2X transcription buffer) and incubating it for 5 min at 37 °C. Transcription was initiated by adding an equal volume of 4X elongation master mix containing 1.2 mM ATP, 1.2 mM GTP, 1.2 mM CTP, 0.4 mM UTP, 0.6 μCi of [α-^32^ P] UTP, 320 mM KCl and 1X transcription buffer (40 mM Tris-HCl, pH 8, 5 mM MgCl_2_, 4 mM dithiothreitol, 0.3 mM EDTA, and 5% glycerol). The reaction mixture was incubated for 25 min at 37 °C and transcription was stopped by adding an equal volume of 2X gel loading buffer (20 mM EDTA, pH 8, 0.025% SDS, 0.025% bromophenol blue, and 0.025% xylene cyanol in formamide). To test the effects of intrinsic termination factors, NusA (4 μM), NusG (4 μM), or both were added to the open complex and incubated for 5 min at 37 °C prior to adding the elongation master mix. DNA oligonucleotides were premixed in 10-, 100-, and 1000-fold molar excess over template with 4X elongation mix and added together when used. The samples were denatured by heating for 8 min at 80 °C and fractionated on standard 5% polyacrylamide sequencing gels. RNA bands were visualized with a phosphoimager and quantified using Image-Quant software (GE Healthcare Life Sciences). Each in vitro transcription experiment was performed twice.

### RNA isolation and RNA sequencing

For total RNA extraction, bacterial cultures were grown in liquid media ON with shaking at 37 °C and diluted 1/100 times in fresh LB media the next day. The fresh cultures were grown at 37 °C with shaking until the culture reached an OD_600_ = 0.8-1, at which time 1.5 mL of the culture was harvested by centrifugation and stored immediately at −80 °C. Pelleted cells were thawed on ice and then lysed using a bead mill. Total RNA was isolated using the Monarch Total RNA Miniprep Kit (NEB) and stored at −80 °C. The concentration of RNA samples was measured using a Nanodrop, and 500 ng of RNA was fractionated through a 1% agarose bleach gel to assess the quality. The RiboCop rRNA depletion kit (Lexogen Vienna, Austria) was used to remove rRNA from 500 ng total RNA. Subsequently, the CORALL Total RNA-SeqTotal RNA Library Kit (Lexogen Vienna, Austria) was used to prepare libraries for Illumina sequencing. Samples were sequenced on the Illumina NextSeq 1000 to generate 100 bases single end reads (100SE) with an average read-depth of 8–12M reads per sample. The quality of the resulting fastq reads were checked using FastQC v0.11.9 (Babraham Bioinformatics, Cambridge) and mapped on the reference genome (*B. subtilis subsp. subtilis* strain 168 (GenBank identifier AL009126.3)) using Bowtie2 v2.4.2 using default settings (46). The resulting SAM files were converted to BAM using SAMtools 1.11, and featuresCounts 2.0.1 was used to obtain the gene counts for each gene annotated in the reference genome taking into account the orientation of genes and reads (47,48). Duplicate samples were summarized by calculating the mean counts for each gene. Bioinformatics analysis of RNA-seq data was done as described previously (22).

### Data availability

RNA-seq data have been deposited to the NCBI SRA repository (Project PRJNA1364995). Flow Cytometry data is available from the corresponding author upon request.

## RESULTS

### A 50 bp region located just upstream of the *conA1* antitermination gene (gene *28*) encodes an intrinsic terminator

A schematic view of the features within the intergenic region between gene *27* (*rco*) and the first gene of the large conjugation operon of plasmid pLS20, gene *28a* (*conAn1*), is shown in Figure 1A. The positions of the conjugation promoter P*_c_*, its transcription start site (TSS), as well as the binding sites for the transcriptional regulator Rco, which is encoded by gene *27* in the opposite direction, were identified previously (23). *conAn1* is preceded by a 456 bp 5’ leader region. The DNA sequence of P*_c_* and the 5’ leader region is shown in Figure 2. Throughout the paper, the positions of the features within the 5’ leader region are numbered relative to the P*_c_* TSS. RNA structure predictions using TransTermHP (transterm.cbcg.umd.edu) (49), as well as Erpin and RNAmotif on the ARNold web server (rna.igmors.u-psud.fr*/*toolbox) (41,42), were performed to predict whether the 5′ leader contains RNA structures. These analyses identified a putative intrinsic terminator (Ter_up28_) just upstream of the Shine-Dalgarno (SD) sequence of *conAn1* (Figures 1 and 2). If this region between positions 389 and 419 of the 5’ leader encodes a terminator, transcripts derived from P*_c_* would terminate upstream of the conjugation genes, resulting in reduced expression of the conjugation operon.

**Figure 2.**
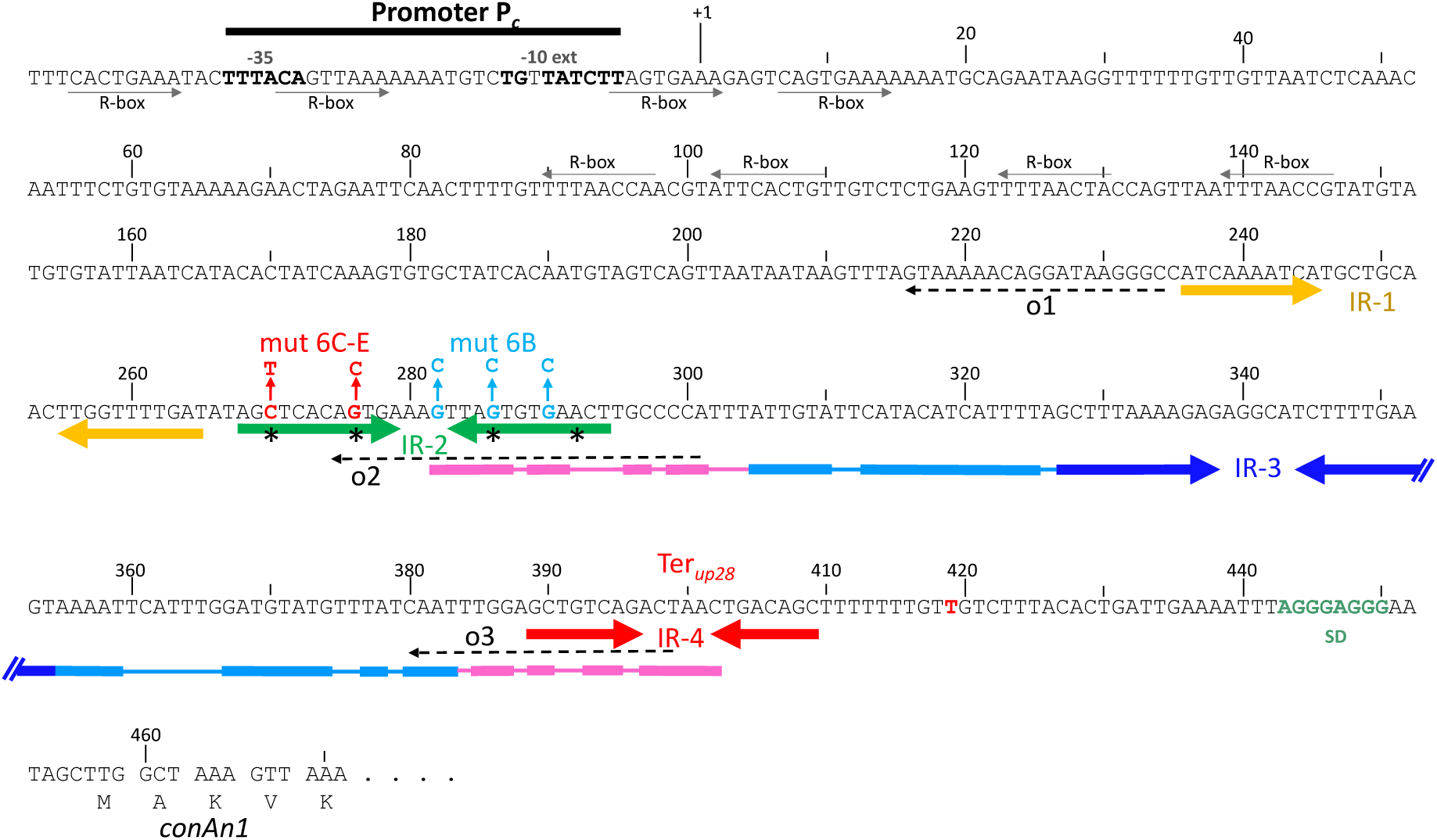
DNA sequence and features of the *conAn1* promoter and leader region. Promoter P*_c_* with an extended −10 sequence, the transcription start site (+1), Rco binding sites (R-box), and the start of gene *28* (*conAn1*) with its SD sequence are indicated. Numbering corresponds to the P*_c_*-derived transcript. The region between positions 230 and 410 contains four inverted repeat sequences indicated with IR-1 (yellow arrows), IR-2 (green arrows), IR-3 (blue arrows) and IR-4 (red arrows). These IRs are predicted to form stem loop structures in the transcript with calculated free energies at 37 °C of −7.5 (SL-1), -6.4 (SL-2), −10.3 (SL-3) and −11.3 (SL-4) kcal/mol. SL-4 is the Ter*_up28_*hairpin. RNA structure predictions suggest that the stem loop derived from IR-3 forms the apex of an extended imperfect stem loop, that can be extended even further to include sequences of SL-2 and Ter_up28_ (see text). These extensions are indicated with light blue and pink lines, respectively. Mismatches and bulges in these regions are indicated with thin lines. The position of the likely transcription termination site for Ter*_up28_*, position 419, is shown in red. Sequences of the three oligonucleotides o1, o2, and o3 used in the *in vitro* transcription experiments are indicated with arrows containing dashed lines.

We used an *in vivo* expression system to test whether this region constitutes a functional terminator. The system, which is based on the *B. subtilis* integration vector pAND101, was used previously to identify and characterize the two-component antitermination system *conAn* of pLS20 (27,28). Briefly, the *amyE* integration cassette present on this vector contains the IPTG-inducible P*_spank_* promoter that drives expression of a *gfp* reporter gene. pAND101 was used to construct the *B. subtilis* control strain AND101 containing a single copy of P*_spank_*-*gfp* integrated at the *amyE* locus. When grown in the absence of IPTG, no GFP was produced in AND101 cells. But when the strain was grown in the presence of IPTG, GFP was produced and cells became fluorescent (27). The fluorescence level of individual cells was measured by flow cytometry. The DNA regions to be tested for termination activity were inserted into this vector between P*_spank_* and *gfp*, generating P*_spank_*-X-*gfp*, where X is the cloned fragment. Each resulting transcriptional fusion was then integrated into the *amyE* locus of the *B. subtilis* chromosome. When grown in the presence of IPTG, the fluorescence levels of cells would be lower than those of the control strain AND101 if the cloned fragment encodes a functional terminator. Several different regions of the 5’ leader were tested. For simplicity, we refer to the cloned fragments using numbers as indicated in Figure 3.

**Figure 3.**
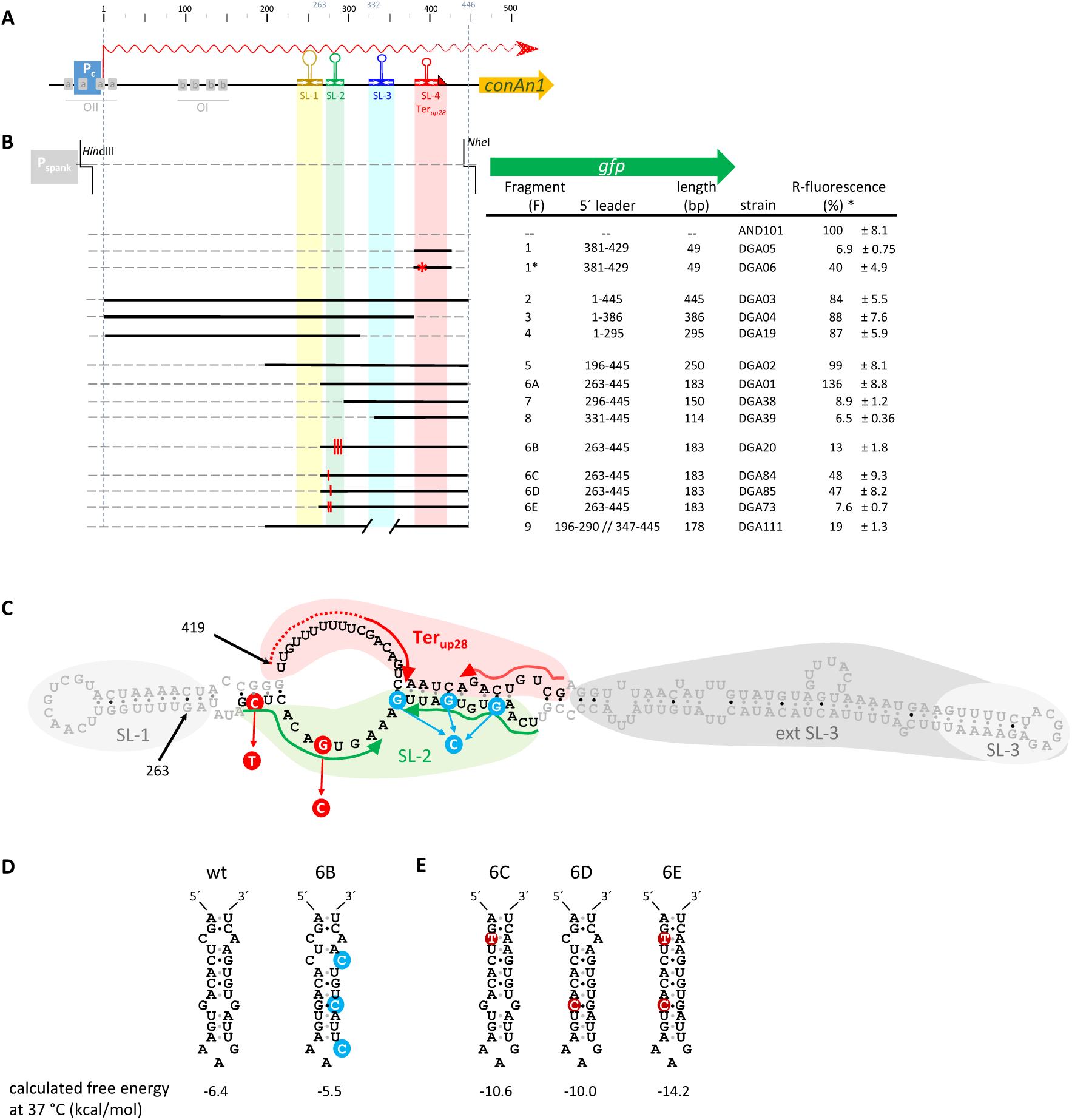
A transcription attenuator precedes the first gene of the pLS20 conjugation operon. **(A)** Schematic diagram of the promoter and leader region of *conAn1*. Inverted repeated sequences 1 to 4, which are predicted to form stem loop structures in the transcript, are indicated with the following colours: yellow (SL-1), green (SL-2), blue (SL-3), and red (SL-4 [Ter*_up28_*]). For other symbols and colors, see Figure 1. **(B)** The top line shows features of the relevant region of AND101 spanning the P*_spank_*promoter and *gfp* (green arrow). The lower panel provides a schematic overview of the different regions of the 5′ leader that are present in the corresponding integrated fusions. Information about the cloned fragments, their name (F), the corresponding strain name, and the fluorescence levels obtained are shown on the right. **(C)** Predicted antitermination conformation of the 5′ leader region spanning positions 231 to 419. Sequences forming SL-1 and the (extended) SL-3 are shown on a grey background. Sequences forming SL-2 and Ter*_up28_*(SL-4) in the predicted termination conformation are highlighted on a green and red background, respectively, and nucleotides that would form the stems of these structures are indicated with arrows. The three Gs that are mutated to Cs in fragment F6B are highlighted in blue. The C and G residues that are mutated in fragments F6C-E are highlighted in red. **(D)** Predicted secondary structures of the wild type (WT) sequence (left) and the triple mutant in fragment F6B (right) of SL-2. G to C mutations are highlighted in blue. **(E)** Predicted secondary structures of SL-2 mutation C270T (fragment F6C, left), G276C (fragment F6D, middle) and the double mutant C270T/G276C (fragment F6E, right). The calculated free energies of these structures are given at the bottom. Note that the mutations in fragment 6C-E, but not the ones in fragment 6B, are predicted to strengthen the secondary structure.

To test if the predicted terminator Ter*_up28_* is functional *in vivo,* a 49 bp region encompassing Ter*_up28_* (positions 381-429, fragment 1 [F1], Figure 3) was cloned upstream of *gfp* in pAND101 to generate pDGA05. In addition, a variant containing a G to A point mutation in the stem of Ter*_up28_* (Fig. 1B) was also constructed (pDGA06). This fragment was named F1*. The two pDGA vectors were used to construct the *B. subtilis* strains DGA05 (P*_spank_*-F1-*gfp*) and DGA06 (P*_spank_*-F1*-*gfp*), which were then analysed for GFP expression together with strain AND101 (negative control) by flow cytometry. In agreement with earlier results (27), the control strain AND101 displayed a high level of GFP fluorescence when grown in the presence of IPTG and this value was set to 100%. Importantly, a low fluorescence level (6.6%) was obtained for strain DGA05, showing that the presence of Ter_up28_ upstream of the *gfp* reporter strongly inhibited expression of *gfp*, implying that the insert encodes a functional terminator. The predicted structure of this terminator suggested that the point mutation present in strain DGA06 would disrupt the 8 bp stem and consequently affect the function of Ter*_up28_*. This prediction was supported by the observation that the fluorescence level obtained for DGA06 was 6-fold higher than that of strain DGA05, which contained the wild type (WT) sequence (Figure 3). Together, these results provide strong evidence that the 49 bp region immediately upstream of the first gene of the conjugation operon (*conAn1, gene 28a*) encodes an intrinsic terminator.

### Ter*_up28_* functions as a transcription attenuator

As Ter*_up28_* is located immediately upstream of the first gene of the conjugation operon, we suspected that this operon would be regulated by a transcription attenuation mechanism in which an alternative antiterminator structure would compete with Ter*_up28_* hairpin formation. Consistent with this hypothesis, our previous RNA-seq results obtained when conjugation levels were at their maximum showed that transcripts starting at P*_c_* bypassed Ter*_up28_* (22,27,50). As a first approach to test this possibility we constructed additional strains, each containing a different fragment of the leader region in front of the *gfp* reporter. As expected, strains DGA04 and DGA19 containing the 5′ leader fragments lacking Ter*_up28_* (fragments F3 and F4), displayed high fluorescence levels (Figure 3). Importantly, high fluorescence levels were also obtained for strain DGA03 containing fragment F2, corresponding to the entire leader region and including Ter*_up28_* (positions 1 to 445, Figures 2 and 3). This result suggested that transcription of the 5’ leader region preceding Ter*_up28_*inhibited terminator function *in vivo*. As explained in more detail below, the 3′ half of the leader transcript contains predicted stem loop structures. We therefore tested whether this RNA segment was important for the putative attenuation mechanism. Strain DGA02 containing *gfp* fused to fragment 5 also displayed a high fluorescence level. These results provided compelling evidence that sequences in the 3′ half of the leader region are responsible for generating an antiterminator structure that competes with formation of Ter*_up28_ in vivo*.

### Formation of Ter*_up28_* is prevented by an antiterminator structure in the 5’ leader

Transcription attenuation mechanisms typically involve the formation of mutually exclusive antiterminator and terminator structures, with the antiterminator sequences preceding and overlapping the terminator. Generally, the conformation that is energetically and/or kinetically more favourable will form by default, while binding of a factor that can compensate for the less favourable structure or interfere with formation of the more favourable structure is required to generate the alternative structure (9). Thus, identifying the default conformation is crucial for understanding the function of the attenuator. We used *in vitro* transcription experiments to examine termination in the 5’ leader. These *in vitro* experiments allowed us to eliminate the contribution of any factor that might bind to the nascent transcript *in vivo* and thereby favour the switch from the default to the alternative structure. A DNA template containing the 3′ half of the leader (positions 199-466) fused to a consensus σ^A^-dependent promoter was used with purified *B. subtilis* RNAP. Transcription termination at Ter*_up28_* occurred at or near position 419. This conclusion is based on the relative migration position of the terminated transcripts compared to migration positions of RNA with known lengths run in parallel (data not shown). We found that the termination efficiency (%T) of Ter*_up28_* was only 3%, indicating that an antiterminator structure was the likely default conformation of the nascent transcript (Figure 4A). We also tested the effect of NusA and NusG on termination because these general transcription elongation factors are known to stimulate intrinsic termination of several hundred intrinsic terminators in *B. subtilis* (43–45). However, neither factor stimulated termination of Ter*_up28_* (Fig. 4A).

**Figure 4.**
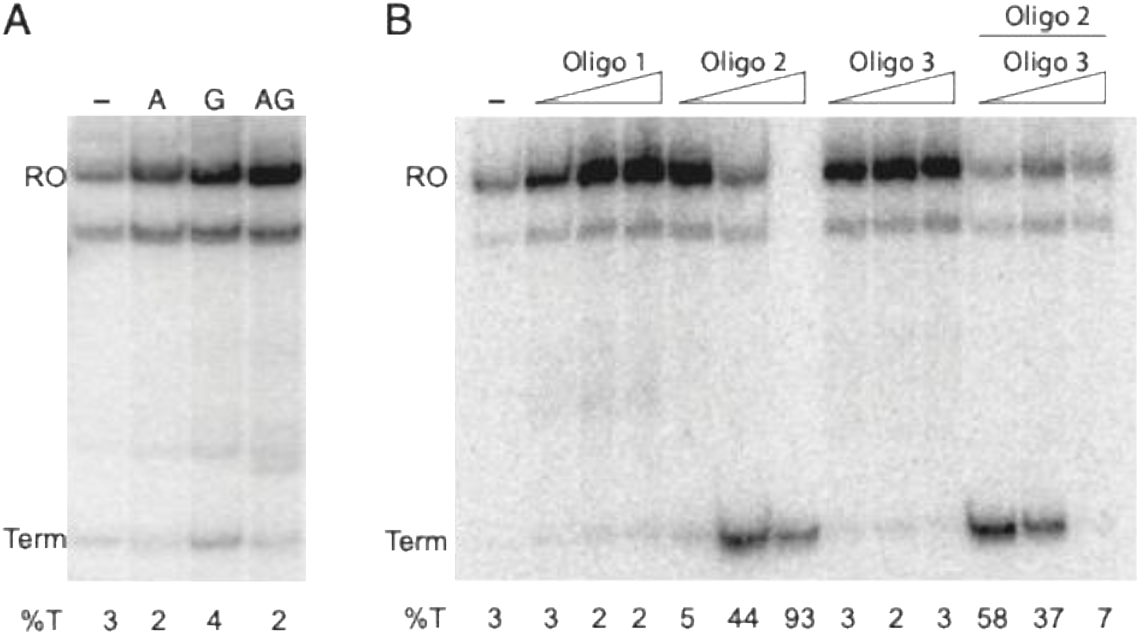
*In vitro* termination assays confirm intrinsic termination and the presence of an antiterminator structure. A DNA fragment containing a consensus σ^A^-dependent promoter sequence followed by positions 199 to 466 of the 5’ leader was used as a template for multi-round *in vitro* transcription assays. (**A**) *In vitro* transcription assay carried out in the absence (–) or presence of NusA (A), NusG (G), or both (AG). Positions of terminated (Term) and run-off (RO) transcripts are marked. Termination efficiency (%T) is shown at the bottom. (**B**) *In vitro* transcription assays performed in the absence (–) or presence of DNA oligonucleotides complementary to 5’ leader positions 216 to 234 (oligo 1), 274 to 301 (oligo 2), or 380 to 399 (oligo 3). Oligos were added in 10-fold, 100-fold, or 1000-fold molar excess over template DNA. In the final three lanes, oligo 2 was held constant at 100-fold molar excess.

### A ⁓20 nt region located 100 nt upstream of Ter*_up28_* is crucial for antitermination

Analysis of the 5’ leader region of gene *28a* revealed that the upstream half does not contain inverted repeats that would be capable of forming stable RNA hairpins. However, four stem-loop (SL) structures were predicted in the downstream half of the leader region, with the Ter*_up28_* hairpin corresponding to SL-4 (positions 389-409) (Figures 2 and 3). This RNA hairpin is relatively stable with a calculated free energy of −11.8 kcal/mol. The other three SLs have free energies of −7.5 (SL-1, positions 236-265), -6.4 (SL-2, positions 268-294), and −10.3 (SL-3, positions 317-352), respectively. These SLs are indicated in Figure 2, and their predicted structures are shown in supplemental Figure S1. To test whether structures corresponding to SL-1, SL-2, or SL-3 can compete with formation of Ter*_up28_*, we constructed three additional *gfp* fusion strains containing successive deletions of fragment F5; fragment 5 was shown above to interfere with Ter*_up28_* function (strain DGA02, Fig. 3). Strain DGA01 (fragment F6A) containing SL-2, SL-3, and SL-4, but not SL-1, displayed high fluorescence levels. In contrast, low fluorescence levels were observed for strain DGA38 (fragment F7) lacking SL-1 and SL-2, and for strain DGA39 lacking SL-1, SL-2, and part of SL-3. These results suggested that SL-2 contains sequences required for antiterminator formation (Figure 3).

The PASIFIC webserver (https://www.weizmann.ac.il/molgen/Sorek/PASIFIC/) is designed to predict alternative RNA structures of putative riboregulators (51). When 5’ leader sequences extending from position 231 to the probable transcription termination site (TTS) at position 419 was examined, a terminator conformation was predicted that included Ter*_up28_* (left panel Fig. 5B). In addition to Ter*_up28_*, SL-1, SL-2, and SL-3 were predicted to form within this structure. To facilitate interpretation, the same colour code used in Figure 3 was used here. Figure 5B also shows that the stem of SL-3 can be extended. The extended version of SL-3 (ext-SL-3) encompasses positions 305 to 383 (indicated with purple ellipse, see also supplemental Fig. S1). PASIFIC also predicted an alternative antiterminator conformation in which the 3′ strand of SL-2 pairs with the 5′ stem and loop sequences of Ter*_up28_* forming a further extended long stem loop, which concomitantly prevents formation of SL-2 and the terminator hairpin (right panel Figure 5B). Remarkably, the calculated minimal free energies of the terminator and the antiterminator conformations were similar, suggesting that little energy is required to switch from one conformation to the other.

**Figure 5.**
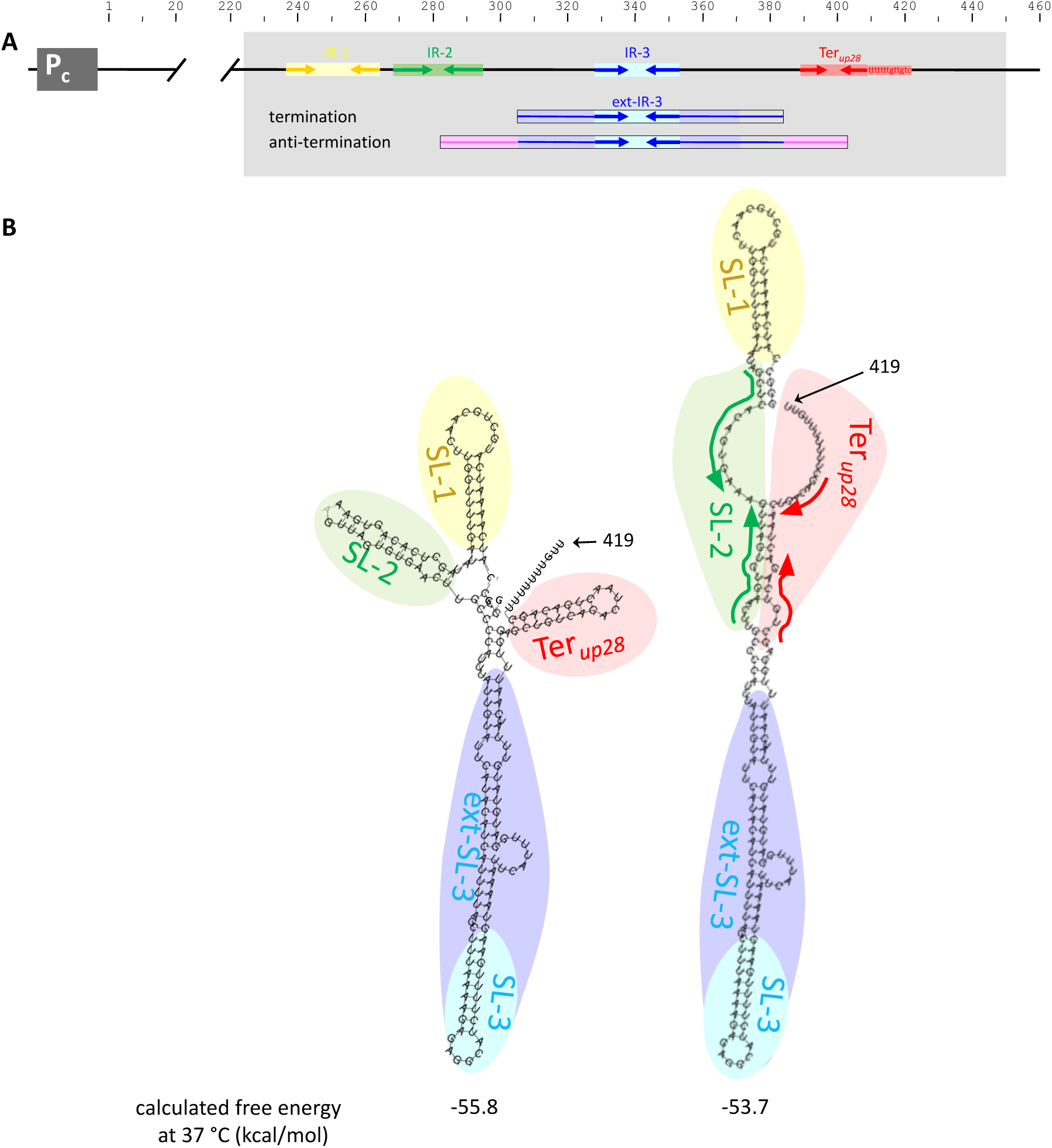
Alternative terminator and antiterminator conformations are predicted to form in the 3′ half of the *conAn1* leader region. **(A)** Schematic view of the P*_c_* promoter and the *conAn1* 5′ leader region. The ⁓ 3′ half of the leader region that contains four stem loops was analysed for predicted secondary structures using the PASIFIC webserver. SL3 forms the apex of a two-stage longer imperfect SL. **(B)** Terminator (left panel) and antiterminator (right panel) conformations predicted for the 5′ leader region spanning positions 231 to 419 by the PASIFIC webserver using “simple antiterminator” settings. Colour codes to visualize the stem loops and the terminator are the same as those used in figures 2 and 3.

The RNAfold webserver (http://rna.tbi.univie.ac.at/cgi-bin/RNAWebSuite/RNAfold.cgi) is designed to predict general local RNA secondary structures (52,53). Essentially the same structures as those predicted by PASIFIC were predicted by the RNAfold webserver when RNA sequences of the 3′ half of the leader region ending either at the probable TTS or 3 nt downstream were analysed (see supplemental Figure S2). Also in this case, similar minimal free energies were predicted for both structures.

Together, the *in silico* analyses suggest that the downstream half of the 5′ leader can adopt two conformations having similar minimal free energies. One of these corresponds to a terminator conformation in which a long central stem loop structure (ext-SL-3) is flanked at the 5′ side by stem loop SL-2 and at the 3′ side by terminator Ter*_up28_*. In the antiterminator conformation, ext-SL-3 is further extended through pairing of SL-2 and Ter*_up28_* sequences, preventing formation of these two stem loops.

### The pLS20 conjugative operon is regulated by a transcription attenuation mechanism

We performed *in vitro* transcription and *in vivo* expression experiments to test our attenuation model. The *in vitro* transcription assays were performed with the same DNA template as described above, in the absence or presence of three DNA oligonucleotides that were complementary to three different RNA segments of the 5’ leader region (see Fig. 2). In each case, experiments were performed with 10-fold, 100-fold, and 1000-fold molar excess of oligo over template DNA. Oligo 1 was complementary to positions 216 to 234, which is just upstream of SL-1. This oligo served as a control because hybridization of the oligo to the nascent RNA would not interfere with the formation of the putative antiterminator structure or Ter*_up28_*. As shown above (Figure 4A), in the absence of any oligo, only about 3% of transcription was terminated, and as expected, the presence of oligo 1 did not affect the termination efficiency (Figure 4B). Oligo 3 was complementary to positions 380-399 such that hybridization would prevent formation of Ter*_up28_*. By itself, this oligo did not affect the already low termination efficiency of Ter*_up28_*. The third oligonucleotide tested, oligo 2, was complementary to positions 274-301. Hybridization of oligo 2 was designed to prevent formation of the predicted antiterminator, thereby promoting formation of the Ter*_up28_* hairpin. Inclusion of this oligo resulted in >90% termination. We also performed experiments with a constant level of oligo 2 (100-fold excess) with increasing concentrations of oligo 3. In this experiment, hybridization of oligo 2 would promote termination by interfering with formation of the antiterminator unless oligo 3 also hybridized to the terminator hairpin. As shown in Figure 4B, oligo 2 promoted termination in the presence of 10-fold and 100-fold excess of oligo 3, but this effect was eliminated at 1000-fold excess of oligo 3 (Figure 4B).

To further test the attenuation model *in vivo*, we introduced mutations into fragment F6A in the context of the *gfp* reporter in pAND101 (Figure 3B) that were predicted to favour formation of Ter*_up28_*. Strain DGA20 contains the *gfp* reporter fusion in which three G to C point mutations were introduced at 5’ leader positions 282, 286, and 290. In the antitermination structure, these Gs anneal with Cs at positions 402, 398, and 394, respectively. The mutations would prevent formation of these base pairs, thereby favouring formation of Ter*_up28_* (Fig. 3C). Note that these three point mutations had only a small effect on the predicted strength of SL-2 (Fig. 3D). Figure 3B shows that GFP expression of this fusion (strain DGA20, fragment 6B) was low, confirming that these mutations favoured formation of Ter*_up28_*as predicted.

According to the attenuation model, the switch from the terminator to the antiterminator conformation is due to the annealing of nucleotides between Ter*_up28_* and SL-2. Consequently, while in the terminator conformation both SL-2 and Ter*_up28_* are formed, formation of these structures is prevented in the antiterminator conformation (Figure 5). SL-2 is weak (calculated free energy of 6.4 kcal/mol) due to two mismatches in its stem. We wondered whether strengthening SL-2 would be sufficient to stimulate formation of Ter*_up28_*. The portion of SL-2 that is predicted to anneal with part of Ter*_up28_* within the antiterminator structure is limited to the 3′ sequence of the stem (Figure 3C). We therefore constructed derivatives of fragment 6A containing single (fragments 6C and 6D) or double point mutations (fragment 6E) in the 5′ portion of the SL-2 stem that would strengthen its structure by removing the mismatches (Figure 3D). Importantly, these substitutions would not interfere with the base pairing that is required for antiterminator formation. These substitutions strengthened SL-2 by about 4 and 8 kcal/mol for the single and double mutations, respectively (Figure 3E). Compared to the WT fragment 6A, expression of *gfp* was reduced by about 65% for the strains containing the single mutations, and it was almost eliminated in the double mutant strain. These results indicate that formation or stabilization of SL-2 favours Ter*_up28_*.

The results from both the *in vitro* and *in vivo* experiments support the following conclusions. First, the 5’ leader adopts the antiterminator conformation by default. Second, formation of the antiterminator is driven by annealing between sequences within SL-2 and Ter*_up28_*. Third, both SL-2 and Ter*_up28_* are concomitantly formed in the terminator conformation. Finally, SL-2 is a weak stem loop and strengthening its secondary structure moderately by a single point mutation causes a substantial fraction of the RNA molecules to adapt the terminator conformation. Taken together, our results demonstrate that transcription attenuation regulates the conjugation operon on pLS20. We name this attenuator cATT*_pLS20_* (conjugation attenuator of pLS20).

### Ext-SL-3 is an essential part of the cATT***_pLS20_*** attenuator

According to the attenuation model, SL-2 and Ter*_up28_*are separated by ⁓ 90 nt that are contained within ext-SL-3 (see Figure 5). As an initial approach to analyse whether this conspicuous feature is important for functionality of the attenuator, we predicted the RNA structure of the 5’ leader region (positions 196-445) lacking the sequences that would form ext-SL-3 (positions 299-370). Using the PASIFIC webserver we predicted that this region would preferentially adopt the terminator conformation (supplemental Figure S3).

To test the importance of the ext-SL-3 region *in vivo*, we constructed an additional reporter strain, DGA111, in which *gfp* is preceded by fragment F9 (see Figure 3B) that contains positions 231-419, but lacking sequences forming ext-SL-3 (positions 299-376). As shown in Figure 3B, fluorescence levels of DGA111 cells were 5-fold lower than those obtained for the parental WT strain DGA02, indicating that ext-SL-3 is important for formation of the antiterminator and proper functioning of cATT*_pLS20_*. In addition to the *gfp* reporter strain DGA111, we tested the effect of deleting the ext-SL-3 region by constructing the pLS20cat derivative pLS20catΔUpTer, in which the leader region spanning positions 299-370 were deleted. The conjugation efficiency of pLS20catΔUpTer was almost 1000-fold lower than that of the WT pLS20cat (3.2E-6 ± 6.3E-7 and 2.7E-3 ± 1.0E-3, respectively). To determine whether this strong decrease in conjugation efficiency was due to altered transcription we used RNA-seq to compare expression of the conjugation operons of plasmids pLS20cat and pLS20catΔUpTer. This deletion resulted in a large decrease in expression of almost all genes in the conjugation operon (Figure 6). The sole exception is gene *61* that is most likely preceded by a strong promoter (unpublished results). Taken together, these results provide compelling evidence that ext-SL-3 plays an important role in proper function of cATT*_pLS20_*.

**Figure 6.**
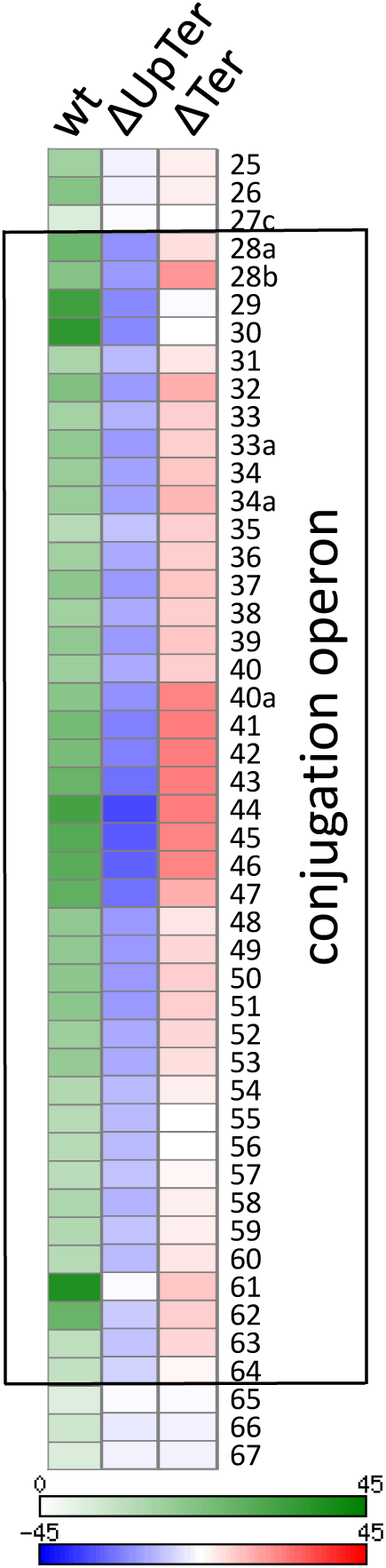
Heat map representation of the expression profiles of the conjugation operons of pLS20cat and derivatives lacking sequences coding for ext-SL-3 or Ter*_up28_*. Left lane (wt) shows the square root of expression levels of genes of the wild type plasmid pLS20cat covering a range from 0 (white, lowest level) to 45 (dark green, highest level). The middle (ΔUpTer) and right (ΔTer) lanes show differential expression levels for these genes of pLS20catΔUpTer and pLS20catΔTer, respectively. Differential expression levels are presented as square root transformation of the normalized coverage on a scale covering a range from −45 to +45 using shades of blue and red for decreased and increased expression, respectively. White reflects no change in expression. Differential expression levels are shown for genes in the conjugation operon genes *28* to *64* (rectangular box) and the three genes upstream and downstream of this operon. Gene numbers according to our nomenclature (22) are given on the right. RNA was isolated from cells harvested in late exponential phase when conjugation efficiency is at its maximum (22).

We also constructed another derivative of pLS20cat, pLS20catΔTer, in which we deleted 51 bp spanning Ter*_up28_*. RNA-seq analysis revealed elevated expression levels of the conjugation genes of pLS20catΔTer (Fig. 6), indicating that in the WT plasmid a portion of the transcripts derived from P*_c_* adopt the terminator configuration. However, despite higher expression levels of the conjugation genes, the conjugation efficiency of pLS20catΔTer was about 30-fold lower than that of the wild type plasmid (8.8E-5 ± 3.5E-6 and 2.7E-3 ± 1.0E-3, respectively). This latter result suggests that expression of the conjugation operon is tightly regulated to achieve maximum conjugation efficiency.

### Conjugation operons present on other plasmids of the pLS20 family are preceded by a cATT_pLS20_-like transcriptional attenuator

We previously showed that pLS20 is the prototype plasmid of a family of conjugative plasmids present in bacilli that share a common genetic organization (34). At the time of publication, this family consisted of 35 plasmids; some had complete plasmid sequences, some were reconstructed partially from individual contigs. Two phylogenetic trees were constructed for these 35 plasmids. One was based on the DNA sequences of the origin of replication regions and the other on protein sequences encoded by eight conjugation and two other conserved genes. The two trees were similar, and according to this analysis these plasmids could be divided into four clades, of which clades 1 (harbouring pLS20) and 3 could be subdivided into three and two subclades, respectively. Screening public databases we identified two additional pLS20 family plasmids, harboured by *Bacillus cabrialesii* strain BIK3 (35) (accession code NZ_JAHKKH010000018) and *B. subtilis* strain NRS6190 (accession code OX419552.1), that both belong to clade 1. Based on established nomenclature (34), we name these plasmids pBcaBIK3 and pBsuNRS6190, respectively.

*In silico* approaches were used to analyse whether the conjugation operons present on other plasmids of the pLS20 family are preceded by a putative transcriptional attenuator. For this endeavor, we extracted the 280 bp region upstream of *conAn1* of each plasmid, and then removed identical sequences, which resulted in 19 unique sequences. A multiple sequence alignment (MSA) showed that there is considerable deviation between these 19 sequences (supplemental Figure S4A). Despite being a short region, the maximum likelihood tree constructed from these sequences (supplemental Figure S4B) was similar to the two previous trees based on the sequences of the origin regions and on ten conserved proteins (34). Subsequent bioinformatics analysis resulted in the identification of a putative intrinsic terminator immediately upstream of the *conAn1* SD sequence in all these plasmids. A MSA limited to the terminator region is shown in Figure 7.

**Figure 7.**
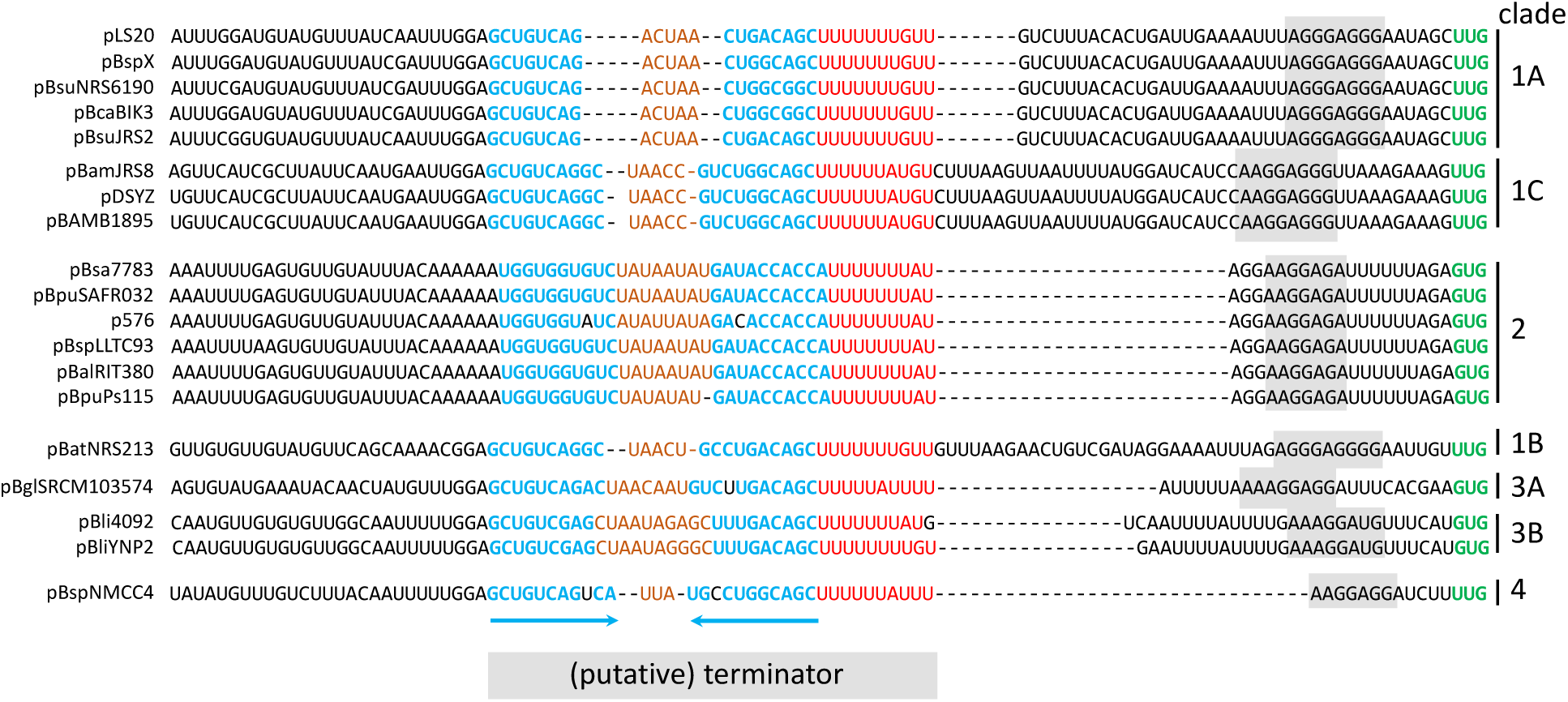
Multiple sequence alignment of the DNA regions encoding the intrinsic terminator upstream of *conAn1* present on pLS20 family plasmids. RNA sequences that would form the stem, loop, and U-rich tract are shown in blue, brown and red colours, respectively. The stem of the terminator is imperfect in three plasmids. These nucleotides are shown in black. The *conAn1* start codon “UUG” or “GUG” is shown in green, and regions containing the SD sequences are highlighted with a grey background. Names of the plasmids and the clade they belong to are given at the left and right of the sequences, respectively.

The MSA of the terminator regions also revealed several other similarities and differences. First, the spacing from the terminator to the SD of the *conAn1* gene varies between 0 and 26 bp. Second, the stem sequences of clade 2 plasmids are considerably different from those of plasmids belonging to the other clades. Finally, an upstream A tract is present in clade 2 plasmids, and the ARNold server predicts that these terminators function as bidirectional terminators.

We then analysed these 19 sequences using the PASIFIC server, which revealed the following. First, RNA regions of all plasmids tested are predicted to adopt a terminator structure that is similar to the one in pLS20 (supplemental Figure S5). Thus, all RNA sequences are predicted to form an ext-SL-3 that is preceded by two stem loops (SL-1 and SL-2) and followed by the terminator stem loop. Second, for 16 out of the 19 sequences, the predicted antiterminator conformation was similar to that of pLS20; i.e. the ext-SL-3 is prolonged through the annealing of sequences of the terminator and SL-2.

The complementary sequences that are involved in antitermination are shown in Supplemental Figure S6. The complementary sequences of the antitermination region are highly conserved for plasmids of clades 1, 3 and 4 (Figure S6). It is worth noting that PASIFIC predicted a distinct antiterminator conformation for plasmid pBatNRS213 of clade 1B, and plasmids pBli4092 and pBliYNP2 of clade 3B. In these three cases, the 5′ strand sequences of the terminator stem are predicted to anneal with 3′ strand sequences of ext-SL-3 rather than with 3′ strand sequences of SL-2 (see Figure S5). However, as shown in Figure S6, the 3′ strand sequences of SL-2 of these three plasmids are nearly identical to those present in other plasmids of clades 1 and 3. This observation suggests that, like the other clade 1, 3 and 4 plasmids, antitermination of these three plasmids are likely driven by annealing of sequences of the terminator and SL-2. Supplemental Figure S6 also shows that the complementary sequences forming the antiterminator region are different for plasmids of clade 2, which is consistent with the observation that the terminator sequence of these plasmids differ from plasmids of the other three clades (see Figure 7). Taken together, these data suggest that a transcriptional attenuator precedes the conjugation operon of all pLS20 family plasmids.

## DISCUSSION

We determined that transcription of the pLS20 conjugation operon is regulated by a transcription attenuation mechanism involving overlapping intrinsic terminator (Ter*_up28_*) and antiterminator structures. This attenuator, which we name cATT_pLS20_, is located in the 5’ leader region preceding the first gene of the conjugation operon, *conAn1*. The downstream half of the RNA leader region can form four stem loops. We show that SL-2, -3 and −4 (SL-4 is the Ter*_up28_* hairpin) participate in the attenuation mechanism (Figures 3 and 4). Under the conditions tested, SL-1 is not essential for cATT_pLS20_ attenuator function. However, it is possible that SL-1 plays a role by acting as an insulator to prevent RNA sequences within the 5′ half of the leader region from interfering with the attenuation mechanism.

*In silico* analysis predicted that SL-3 forms the apex of the longer ext-SL-3. In the terminator conformation, SL-2 and Ter*_up28_* are located near the 5′ and 3′ of the base of ext-SL-3, respectively (Figure 5). Sequences of the 3′ stem of SL-2 and the 5′ stem and loop of Ter*_up28_* are complementary allowing the already long stem of ext-SL-3 to be extended further, thereby adopting the antiterminator conformation while simultaneously preventing formation of SL-2 and Ter*_up28_* (Figure 5). Thus, formation of the antiterminator occurs via a zipper-type mechanism that begins with ext-SL-3, which is propagated until the antiterminator structure is fully formed. Consequently, ext-SL-3 is an important component of the attenuator. This view is supported by the following results. First, our *in vitro* transcription studies showed that the antiterminator structure forms by default and that disruption of the antiterminator with an oligonucleotide promotes termination (Figure 4). Second, instead of the antiterminator, the terminator structure is formed in the absence of ext-SL-3 (Figures 3, 6, and S3). Numerous transcription attenuation mechanisms have been described, but we believe this is the first example of a zipper-type attenuator.

SL-2 has a pivotal role in the switch from the antiterminator to the terminator structure. SL-2 is the weakest of the four stem loops (-6.4 kcal/mol), suggesting that its structure is dynamic and easily destabilized. Mutations to SL-2 that eliminated one or both of the two mismatches in the stem strengthened the structure by −4 to -8 kcal/mol, and this caused a large decrease in expression, consistent with the mutations causing most or all of the transcripts to adopt the terminator conformation (Fig. 3). These results show that switching between the antitermination and the termination conformations requires little energy, which is consistent with the similar calculated free energies for the terminator and antiterminator conformations (Figure 5). SL-2 is located upstream of Ter*_up28_* and hence the SL-2 sequences are synthesized before those of Ter*_28up_*. Thus, formation and/or stabilisation of SL-2 promotes formation of the terminator because it prevents completion of the antiterminator. Interestingly, the MSA presented in Figure S4 shows that the SL-2 region, and particularly the 5′ stem sequence, is the most conserved region of the 19 leader sequences of pLS20 family plasmids. This observation suggests that in addition to its structure, the RNA sequence may serve another important function by recruiting an RNA binding protein that might stabilize SL-2, and thereby promote terminator formation. While we have tested several possible RNA or DNA binding proteins encoded by pLS20 (Rco, ConAn1, p34a, p35, p74A, p74B, p74C, p64), we did not find evidence for any of these playing a role in the attenuation mechanism. Alternatively, it is possible that a counter transcript that interferes with antiterminator formation is responsible for promoting termination. This type of counter transcript mechanism is responsible for inducing termination in the attenuator region that controls expression of the conjugation operon in the *Enterococcus faecalis* plasmid pCF10 (54).

We previously showed that pLS20 is the archetype of a family of related plasmids that all contain a long conjugation operon that is regulated by a *conAn*-type P-AT system (34). Bioinformatics analysis indicated that all pLS20 family plasmids contain a zipper-type transcriptional attenuator upstream of their *conAn1* gene. In addition to the pLS20 family plasmids, a *conAn*-type P-AT system is located at the start of the conjugation operon on >350 plasmids of Gram-positive bacteria (27). A similar strategy that was used for the pLS20 family plasmids was employed to determine whether these plasmids also contain a putative attenuator preceding their conjugation operon. By screening the plasmid database (39,40), we identified 1,162 plasmids containing a *conAn1* homolog followed by typical conjugation genes such as ATPase encoding genes *virB4* and *virD4*. After removing identical sequences and those corresponding to pLS20 family plasmids, 339 of the remaining 423 unique sequences were predicted to contain a terminator, which in most cases was located just upstream of the *conAn1* gene. Additional analyses suggested that these terminators form part of a transcriptional attenuator, and in many cases these attenuators appear to be of the zipper-type, which is a particularly common arrangement for attenuators present on plasmids from lacticaseibacilli.

Our results from this and prior studies provide the following model of how numerous conjugation operons present on plasmids from Gram-positive bacteria are regulated. Expression of the conjugation operons is repressed by default, and a complex multi-tiered regulatory system enables activation of the conjugation process under suitable conditions. This multi-tiered system includes at least three distinct levels. The first level regulates transcription initiation via a quorum sensing signal peptide system that controls the activity of the main conjugation promoter through derepression. The other two levels act during transcription elongation. The first of these two layers comprises the zipper-type attenuator described here. At least in pLS20, the attenuator adopts the antiterminator conformation by default permitting transcription to proceed. However, the attenuator can function as a check-point mechanism that may terminate transcription of the conjugation genes. There are two non-mutually exclusive circumstances when this may occur. First, it may act as a sensing mechanism that occurs under conditions not suitable or compatible with the conjugation process. Second, it may act as a negative feedback loop to prevent overexpression of the conjugation genes. This latter scenario is supported by the observation that the conjugation genes are overexpressed in the absence of cATT_pLS20_ and that overexpression leads to a conjugation defect. Future studies are needed to analyze these possibilities. The second layer acting during transcription elongation is the *conAn*-type P-AT system. Contrary to the zipper-type attenuator that acts locally by controlling a single terminator, the P-AT system allows the transcription elongation complex to bypass multiple intrinsic terminators. In addition to regulating transcription processivity, the P-AT system minimizes the effects of spurious transcription and allows differential expression of subsets of genes within the conjugation operon (27). Thus, tight control of conjugation is achieved to prevent this energetically costly process from occurring in the absence of appropriate recipient cells and under otherwise unfavorable conditions.

## ACKNOWLEDGEMENTS

We thank Harma Spakman for technical help with RNA-seq and members of the laboratory for fruitful discussions. The funders had no role in study design, data collection and analysis, decision to publish, or preparation of the manuscript.

## AUTHOR CONTRIBUTIONS

Daniel González Álvarez: Formal analysis, Methodology, Writing—original. Andrés Miguel-Arribas, Conceptualization, Formal analysis, Methodology, Writing—original draft. Sandeepani Ranaweera: Formal analysis, Methodology. Fernando Freire-Gómez: Writing—original, Methodology. David Abia: Conceptualization, Methodology, Visualization. Anne de Jong: Formal analysis, Methodology, Ling Juan Wu: Methodology, Validation, Writing—original draft. Paul Babitzke, Conceptualization, Formal analysis, Methodology, Validation, Writing—original draft, Writing—review & editing. Wilfried Meijer: Conceptualization, Formal analysis, Methodology, Validation, Writing—original draft, Writing—review & editing.

## SUPPLEMENTARY DATA

Supplementary Data are available at NAR online.

## DATA AVAILABILITY

RNA-seq data have been deposited to the NCBI SRA repository Project PRJNA1364995. Flow Cytometry data is available from the corresponding author upon request. All other relevant data are given in the main text.

