## Supplemental data for "EXPRESSION OF CONJUGATION GENES IS CONTROLLED BY PROCESSIVE ANTITERMINATION AND A NOVEL ZIPPER-TYPE TRANSCRIPTIONAL ATTENUATION MECHANISM"

By González Álvarez et al.

### Supplemental Figures

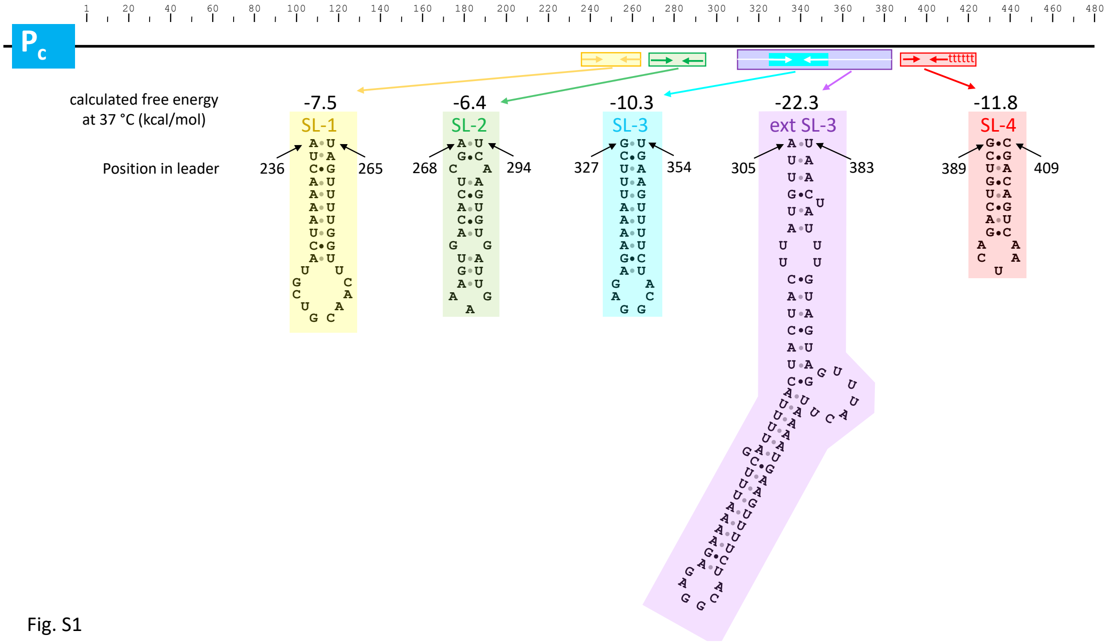

Fig. S1

**Figure S1. The downstream half of the *conAn1* leader contains several RNA structures.** Promoter $P_c$ is shown at the top followed by the 5' leader region and the positions of the stem loop structures. The predicted RNA secondary structures predicted using the Vienna RNAfold (http://rna.tbi.univie.ac.at/cgi-bin/RNAWebSuite/RNAfold.cgi) using default settings are shown at the bottom. The positions within the leader region and the calculated minimum free energy of each structure are also shown.

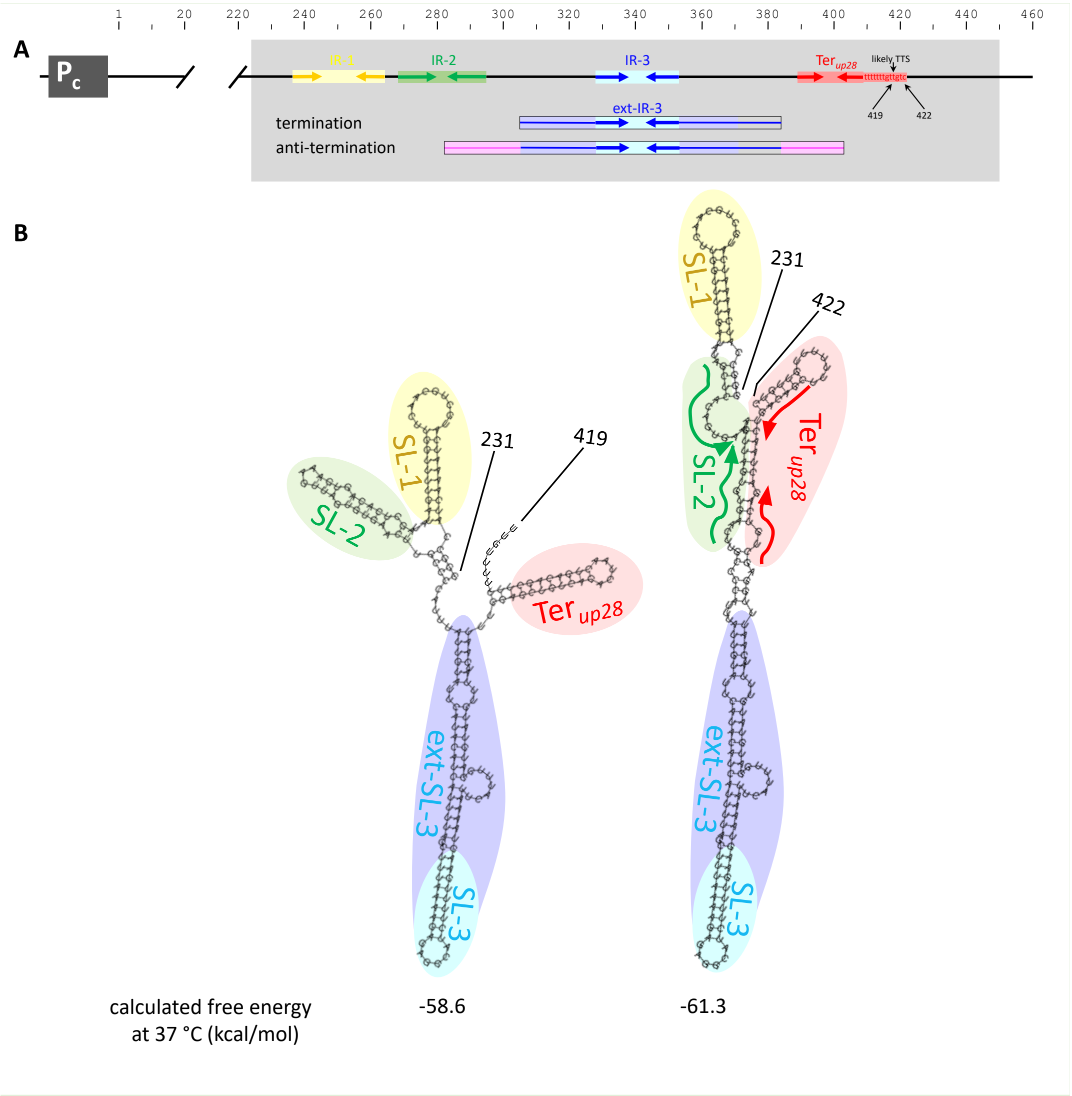

**Supplemental Figure S2.** Alternative terminator and antiterminator conformations predicted in the 3´ half of the *conAn1* leader region by RNAfold. **(A)** Schematic view of the $P_c$ promoter and the *conAn1* 5´ leader region. The ~ 3´ half of the leader region that contains four stem loops was analysed for predicted secondary structures using the RNAfold webserver with default settings. SL3 forms the apex of a two-stage longer imperfect SL. **(B)** Depending on the 3´ end point, the RNA is predicted to adopt a terminator or antiterminator conformation. A terminator conformation was predicted for the leader region spanning positions 231 to 419 (left panel). On the other hand, an antiterminator conformation was predicted for a region spanning positions 231 to 422, or any position further downstream (left panel). The calculated free energy at 37 °C is given below each predicted structure. Colour codes to visualize the stem loops and the terminator are the same as those used in figures 2 and 3.

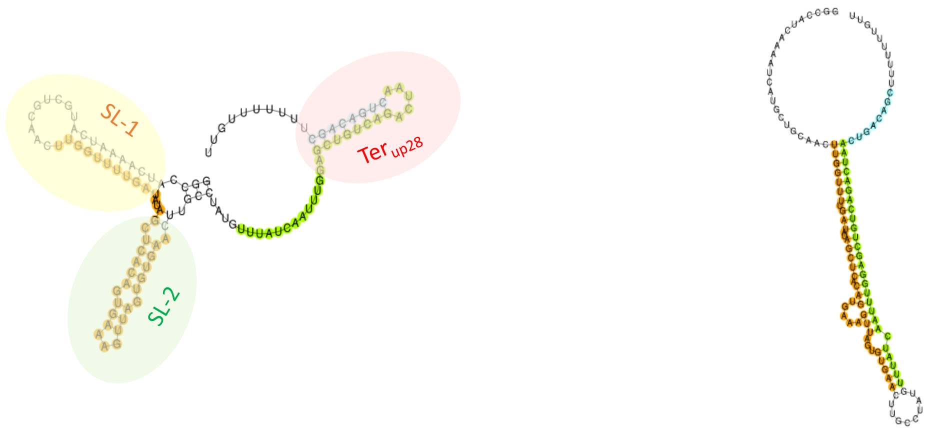

calculated free energy at 37 °C (kcal/mol) -30.0 -15.8

**Supplemental Figure S3.** The terminator configuration is predicted for cATT*pLS20* lacking ext-SL-3 sequences. Terminator and antiterminator structure of the downstream half of the pLS20 5´ leader lacking the region corresponding to ext-SL-3 (positions 231-298/371-419) were predicted with the PASIFIC webserver (https://www.weizmann.ac.il/molgen/Sorek/PASIFIC/). Note the large difference in calculated free energies between the predicted terminator and antiterminator structure (-30 and -15.8 kcal/mol). A low probability score (0.14) was predicted for the antitermination structure.

A

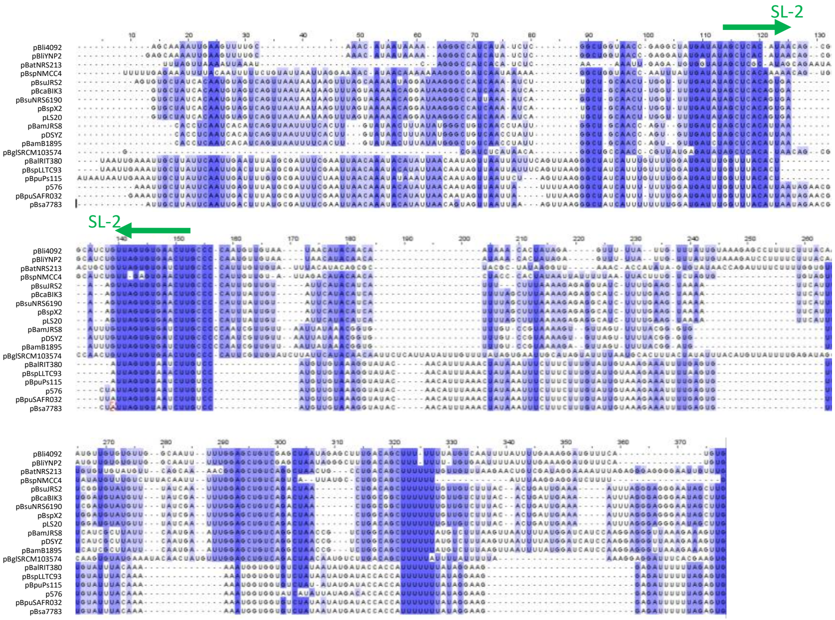

B

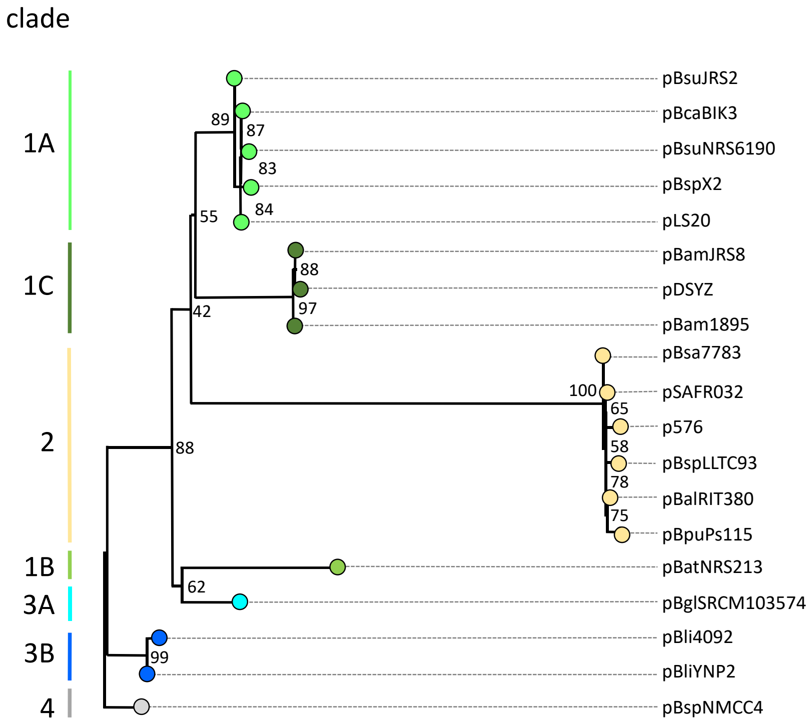

**Supplemental Figure S4.** Phylogenetic conservation of the attenuator identified in pLS20. **A.** Multiple sequence alignment (MSA) of 280 nt of the conjugative operon 5' leader region of 19 unique sequences of pLS20 family plasmids. The MSA was generated with the program LocARNA

(36) using default settings. Level of conservation is indicated with different shadings of blue backgrounds; residues that are >80% conserved are shown on a dark blue background. Dots reflect introduced gaps. Nucleotides forming the stem portions of SL-2 are indicated with a green arrow. **B.** Phylogenetic relationship of the 5' leader regions of 19 different plasmids of the pLS20 family inferred from maximum likelihoods. Statistical evidence for each branch is provided by bootstrap analysis (1000 replicates indicated as percentages). The roots are located at the midpoint. Previously, pLS20 family plasmids were grouped into four clades of which clades 1 and 3 were divided into 3 and 2 subclades, respectively (34). The clade of the plasmids used for the construction of this tree is indicated on the right, as well as with colours: clade 1, green; clade 2, yellow; clade 3, blue; clade 4, grey. Dendroscope (55) version 3.6.3 was used to visualize the tree. For clarity, a final output was generated with power point in which coloured circles were used to label the (sub)clades.

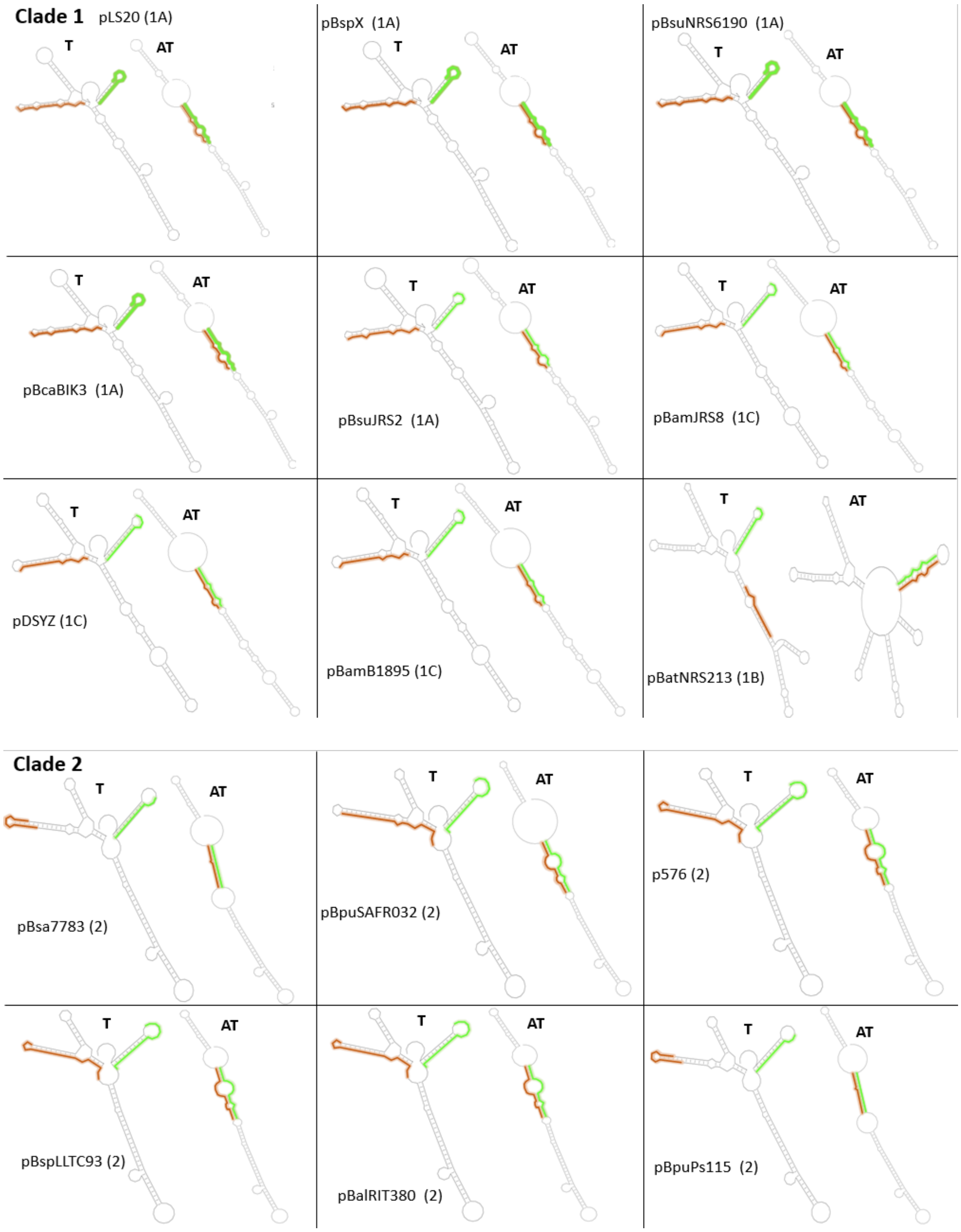
Clade 1
pLS20 (1A)
T
AT
pBspX (1A)
T
AT
pBsuNRS6190 (1A)
T
AT
T
AT
pBcaBIK3 (1A)
T
AT
pBsuJRS2 (1A)
T
AT
pBamJRS8 (1C)
T
AT
pDSYZ (1C)
T
AT
pBamB1895 (1C)
T
AT
pBatNRS213 (1B)
Clade 2
T
AT
pBsa7783 (2)
T
AT
pBpuSAFR032 (2)
T
AT
p576 (2)
T
AT
pBspLLTC93 (2)
T
AT
pBalRIT380 (2)
T
AT
pBpuPs115 (2)

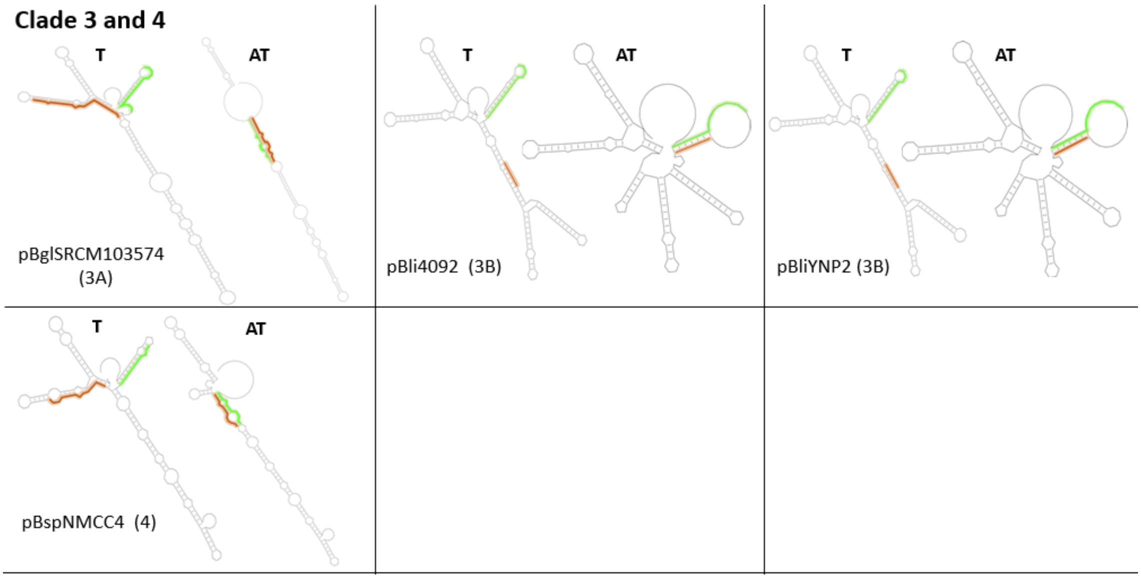

**Supplemental Figure S5.** Terminator and antiterminator conformations of the 5´ leader regions of pLS20 family members. PASIFIC was used to predict the terminator (T) and antiterminator conformation (AT) for 19 unique 5' leader regions belonging to pLS20 family plasmids. Sequences of the 5´ strand of the terminator stem that were predicted to base pair with sequences in the antiterminator are indicated with green and orange lines, respectively. The (sub)clade for plasmid is in brackets following the name of the plasmid. The 3´ ends of the sequences correspond to the transcription termination site identified for pLS20. The 5´ ends correspond to the conserved motif GGGCBRTCA located immediately upstream of SL-1 (B, not A; R, A or G).

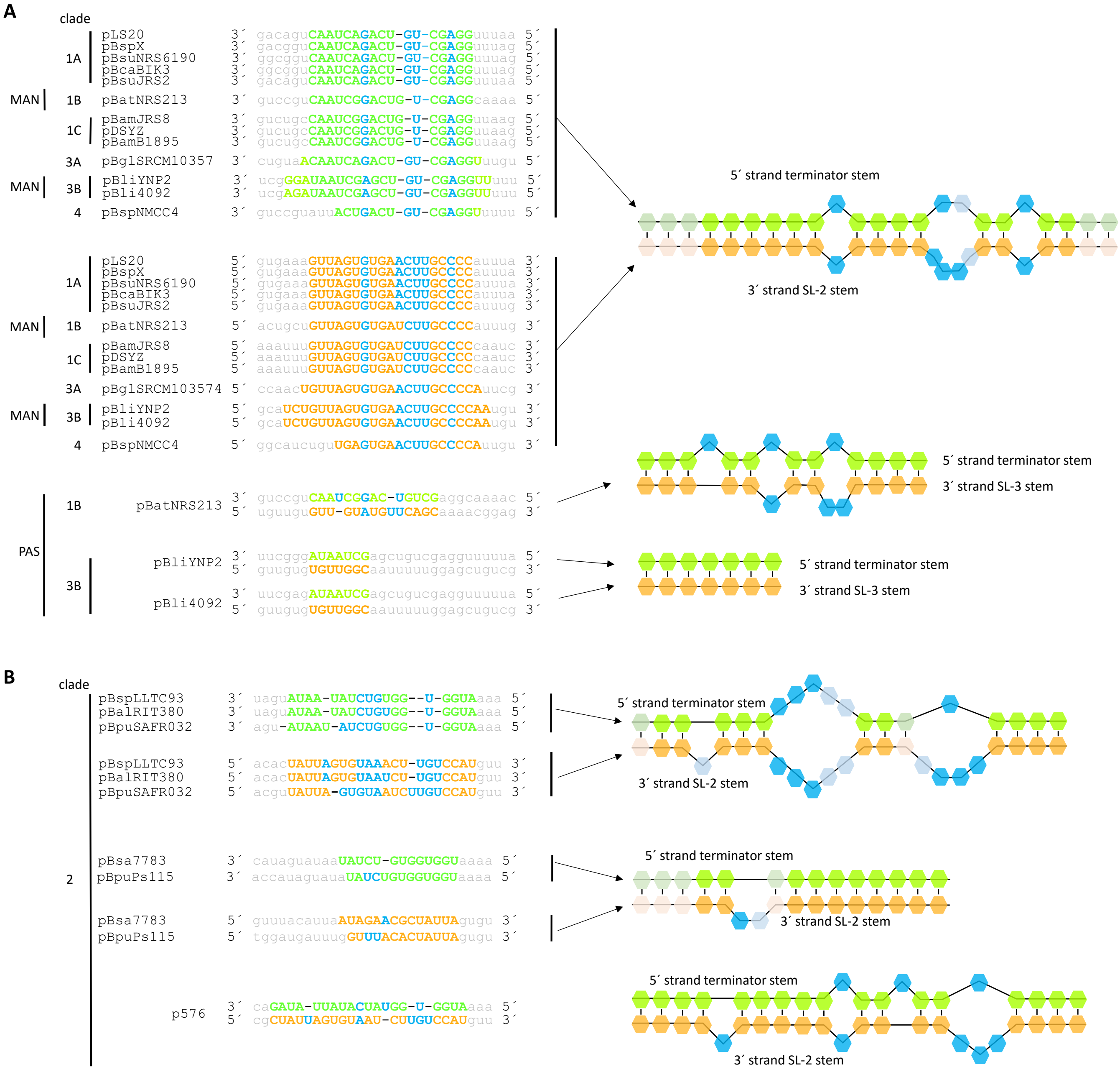

**Supplemental Figure S6.** Complementary sequences in the antiterminator of pLS20 family plasmids. The sequences predicted by PASIFIC to base pair in the antiterminator (AT) are shown for unique sequences of plasmids belonging to clade 1, 3, 4 (panel A) and clade 2 (panel B) of pLS20 family plasmids. The same colour code as that in supplemental Figure S5 is used here: green and orange correspond to sequences that form part of the alternative terminator and SL-2 structures. Non-complementary nucleotides are indicated in blue. For three plasmids, pBatNRS213 (clade 1B), pBli4092 (clade 3B), and pBliYNP2 (clade 3B), the terminator sequences were predicted to base pair with sequences in ext-SL-3, instead of SL-2 (PAS). However, manual inspection revealed that a portion of SL-2 is also capable of base pairing with the terminator sequence as for the other plasmids (MAN). Schematic view of base pairing is shown at the right.

### Supplemental Tables

| Supplemental Table S1. Strains used | | |
|---|---|---|
| **Strains** | **Description and genotype** | **Source or Reference** |
| *Escherichia coli* | | |
| XL1-Blue | Used for regular cloning. *end*A1 *gyr*A96(nal$^R$) *thi*-1 *rec*A1 *rel*A1 *lac* *gln*V44 F'[ ::Tn10 *pro*AB$^+$ *lac*I$^q$ Δ(*lac*Z)M15] *hsd*R17 ($r_K^-$ $m_K^+$) | Laboratory stock. Stratagene |
| JM101 | Used for pMiniMad protocol. *gln*V44 *thi*-1Δ(*lac-pro*AB) F′[*lac*I$^q$ZΔM15 *tra*D36 *pro*AB+] (*rec*A$^+$, $r_k^+$) | Laboratory stock. NEB |
| *Bacillus subtilis* | | |
| 168 (1A700) | *trpC2* | BGSC* |
| PKS11 | *trpC2,* pLS20cat (Cm$^R$) | (Singh et al. 2013) |
| AND81 | *trpC2,* pLS20Δ28 (Cm$^R$) | (Miguel-Arribas et al. 2021) |
| AND101 | *trpC2*, *amy*E::P$_{spank}$-*gfp* (Spec$^R$) | (Miguel-Arribas et al. 2021) |
| DGA05 | *B. subtillis 168* competent cells transformed with the plasmid pDGA05. *trpC2*, *amy*E::P$_{spank}$-F1-*gfp* (Spec$^R$) | This work |
| DGA06 | *B. subtillis 168* competent cells transformed with the plasmid pDGA06. *trpC2*, *amy*E::P$_{spank}$-F1*-*gfp* (Spec$^R$) | This work |
| DGA03 | *B. subtillis 168* competent cells transformed with the plasmid pDGA03. *trpC2*, *amy*E::P$_{spank}$-F2-*gfp* (Spec$^R$) | This work |
| DGA04 | *B. subtillis 168* competent cells transformed with the plasmid pDGA04. *trpC2*, *amy*E::P$_{spank}$-F3-*gfp* (Spec$^R$) | This work |
| DGA19 | *B. subtillis 168* competent cells transformed with the plasmid pDGA07. *trpC2*, *amy*E::P$_{spank}$-F4-*gfp* (Spec$^R$) | This work |
| DGA02 | *B. subtillis 168* competent cells transformed with the plasmid pDGA02. *trpC2*, *amy*E::P$_{spank}$-F5-*gfp* (Spec$^R$) | This work |
| DGA01 | *B. subtillis 168* competent cells transformed with the plasmid pDGA01. *trpC2*, *amy*E::P$_{spank}$-F6A-*gfp* (Spec$^R$) | This work |
| DGA38 | *B. subtillis 168* competent cells transformed with the plasmid pDGA17. *trpC2*, *amy*E::P$_{spank}$-F7-*gfp* (Spec$^R$) | This work |
| DGA39 | *B. subtillis 168* competent cells transformed with the plasmid pDGA18. *trpC2*, *amy*E::P$_{spank}$-F8-*gfp* (Spec$^R$) | This work |

| | | |
|---|---|---|
| DGA20 | *B. subtillis 168* competent cells transformed with the plasmid pDGA08. *trpC2*, *amy*E::$P_{spank}$-F6B-*gfp* ($Spec^R$) | This work |
| DGA84 | *B. subtillis 168* competent cells transformed with the plasmid pDGA67. *trpC2*, *amy*E::$P_{spank}$-F6C-*gfp* ($Spec^R$) | This work |
| DGA85 | *B. subtillis 168* competent cells transformed with the plasmid pDGA68. *trpC2*, *amy*E::$P_{spank}$-F6D-*gfp* ($Spec^R$) | This work |
| DGA73 | *B. subtillis 168* competent cells transformed with the plasmid pDGA66. *trpC2*, *amy*E::$P_{spank}$-F6E-*gfp* ($Spec^R$) | This work |
| DGA111 | *B. subtillis 168* competent cells transformed with the plasmid pDGA92. *trpC2*, *amy*E::$P_{spank}$-F9-*gfp* ($Spec^R$) | This work |
| DGA41 | *trpC2,* pLS20ΔUpTer ($Cm^R$) | This work |
| DGA42 | *trpC2,* pLS20ΔTer ($Cm^R$) | This work |

*, BGSC: *Bacillus* Genetic Stock Center, Department of Bioche*m*istry, The Ohio State University, Columbus, OH, USA. (http://www.bgsc.org/.)

| **Supplemental Table S2. Plasmids used** | | |
|---|---|---|
| **Plasmid** | **Description** | **Source or Reference** |
| pLS20cat | Native plasmid pLS20 labelled with Cm resistance cassette in the unique *Sal*I site. ($Cm^R$) | Itaya et al.,2006 |
| pMiniMAD2 | Plasmid used for marker-less deletions. ($Amp^R$) and ($Em^R$). | gift of Daniel Kearns |
| pAND101 | pDR110 derivate containing promoter less *sfGFP* gene (present in pKsfGFP). ($Amp^R$) and ($Spec^R$). | (Miguel-Arribas et al. 2021) |
| pDGA05 | pAND101 derivate containing fragment F1. Cloned fragment was made by hybridization of primers oDG01 (*Sal*I) and oDG02 (*Nhe*I). ($Amp^R$) and ($Spec^R$). | This work |
| pDGA06 | pAND101 derivate containing fragment F1*. Cloned fragment was made by hybridization of primers oDG01 (*Sal*I) and oDG02 (*Nhe*I). ($Amp^R$) and ($Spec^R$). | This work |
| pDGA03 | pAND101 derivate containing fragment F2. Cloned fragment was amplified using primers oDG05 (*Hind*III) and oDG06 (*Nhe*I). ($Amp^R$) and ($Spec^R$). | This work |
| pDGA04 | pAND101 derivate containing fragment F3. Cloned fragment was amplified using primers oDG05 (*Hind*III) and oDG09 (*Nhe*I). ($Amp^R$) and ($Spec^R$). | This work |
| pDGA07 | pAND101 derivate containing fragment F4. Cloned fragment was amplified using primers oDG05 (*Hind*III) and oDG10 (*Nhe*I). ($Amp^R$) and ($Spec^R$). | This work |
| pDGA02 | pAND101 derivate containing fragment F5. Cloned fragment was amplified using primers oDG04 (*Hind*III) and oDG06 (*Nhe*I). ($Amp^R$) and ($Spec^R$). | This work |
| pDGA01 | pAND101 derivate containing fragment F6A. Cloned fragment was amplified using primers oDG03 (*Hind*III) and oDG06 (*Nhe*I). ($Amp^R$) and ($Spec^R$). | This work |
| pDGA17 | pAND101 derivate containing fragment F7. Cloned fragment was amplified using primers oDG23 (*Hind*III) and oDG06 (*Nhe*I). ($Amp^R$) and ($Spec^R$). | This work |
| pDGA18 | pAND101 derivate containing fragment F8. Cloned fragment was amplified using primers oDG24 (*Hind*III) and oDG06 (*Nhe*I). ($Amp^R$) and ($Spec^R$). | This work |
| pDGA08 | pAND101 derivate containing fragment F6B. Cloned fragment was amplified using primers oDG18B (*Hind*III) and oDG06 (*Nhe*I). ($Amp^R$) and ($Spec^R$). | This work |
| pDGA67 | pAND101 derivate containing fragment F6C. Cloned fragment was amplified using primers oDG88 (*Hind*III) and oDG06 (*Nhe*I). ($Amp^R$) and ($Spec^R$). | This work |

| | | |
|---|---|---|
| pDGA68 | pAND101 derivate containing fragment F6D. Cloned fragment was amplified using primers oDG127 (*Hind*III) and oDG06 (*Nhe*I). ($Amp^R$) and ($Spec^R$). | This work |
| pDGA66 | pAND101 derivate containing fragment F6E. Cloned fragment was amplified using primers oDG85 (*Hind*III) and oDG86 (*Nhe*I). ($Amp^R$) and ($Spec^R$). | This work |
| pDGA92 | pAND101 derivate containing fragment F9. Cloned fragment was amplified using primers oDG146 (*Hind*III) and oDG147 (*Nhe*I), the template was syntetized as gBlock54 by IDT. ($Amp^R$) and ($Spec^R$). | This work |
| pDGA12 | pMiniMAD2 derivate to create pLS20ΔUpTer (DGA41 strain). Cloned fragment was amplified by overlapping PCR using primers oDG11 (*Hind*III) and oRG17 (*Bam*HI). Overlapped fragments “Up” and “Down” were amplified using oDG11/oDG12 and oDG15/oRG17 respectively. ($Amp^R$) and ($Em^R$) | This work |
| pDGA16 | pMiniMAD2 derivate to create pLS20ΔTer (DGA42 strain). Cloned fragment was amplified by overlapping PCR using primers oDG11 (*Hind*III) and oRG17 (*Bam*HI). Overlapped fragments “Up” and “Down” were amplified using oDG11/oDG13 and oDG14B/oDG17 respectively. ($Amp^R$) and ($Em^R$) | This work |
| pLS20ΔUpTer | Deletion of intergenic region of UpTer region on plasmid pLS20cat.) | This work |
| pLS20ΔTer | Deletion of intergenic region of Ter region on plasmid pLS20cat | This work |

**Supplemental Table S3. Oligonucleotides used**

| Name | Sequence (5’-3’) | Purpose |
|---|---|---|
| pDR111_U_sec | TGACTTTATCTACAAGGTGTGGC | Forward primer to verify sequence of PCR fragments cloned in pAND101. |
| pDR111_L_sec | TTAAATGCAACCGTTTTTTCGGAAGG | Reverse primer to verify sequence of PCR fragments cloned in pAND101. |
| oSeqpKSGFP_Dn | TTCCGGCATGGCGGACTTGAAGAAGTC | Reverse primer to verify sequence of PCR fragments cloned in pAND101. |
| oM13_Fw_21_ext | ACGTTGTAAAACGACGGCCAGTG | Forward primer to verify sequence of PCR fragments cloned in pMiniMad2 |
| oM13_Rev_Ext | CACAGGAAACAGCTATGACCATG | Reverse primer to verify sequence of PCR fragments cloned in pMiniMad2 |
| oDG01 | **AGCT**AATTTGGAGCTGTCAGACTAACTGACAGCTTTTTTTGTTGTCTTTACAC | Hybridization primer to make fragment F1. Used in combination with oDG02. *Nhe*I restriction site extension |
| oDG02 | **CTAG**GTGTAAAGACAACAAAAAAAGCTGTCAGTTAGTCTGACAGCTCCAAATT | Hybridization primer to make fragment F1. Used in combination with oDG01. *Nhe*I restriction site extension |
| oDG05 | **TTTTAAGCTT**AAGAGTCAGTGAAAAAAATGCAGAATAAGG | Forward primer to amplify fragment F2, F3 and F4. Used in combination with primer oDG06, oDG09 and oDG10. *Hind*III restriction site extension |
| oDG06 | **TTTTGCTAGC**CCTAAATTTTCAATCAGTGTAAAGACAAC | Reverse primer to amplify fragment F2, F5, F6A, F7, F8, F6B, F6C and F6D. Used in combination with primer oDG05, oDG04, oDG03, oDG23, oDG24, oDG18B, oDG88 and oDG127. *Nhe*I restriction site extension |
| oDG09 | **TTTTGCTAGC**CAAATTGATAAACATACATCCA | Reverse primer to amplify fragment F3. Used in combination with primer oDG05. *Nhe*I restriction site extension |
| oDG10 | **TTTTGCTAGC**AAGTTCACACTAACTTTCACTGTGAGCT | Reverse primer to amplify fragment F4. Used in combination with primer oDG05. *Nhe*I restriction site extension |
| oDG04 | **TTTTAAGCTT**GTCAGTTAATAATAAGTTTAGTAAAAACAGG | Forward primer to amplify fragment F5. Used in combination with primer oDG06. *Hind*III restriction site extension |
| oDG03 | **TTTTAAGCTT**GATATAGCTCACAGTGAAAGTTAGTGTG | Forward primer to amplify fragment F6A. Used in combination with primer oDG06. *Hind*III restriction site extension |
| oDG23 | **TTTTAAGCTT**GCCCCATTTATTGTATTCATACATCATTTTAGCTTTA | Forward primer to amplify fragment F7. Used in combination with primer oDG06. *Hind*III restriction site extension |
| oDG24 | **TTTTAAGCTT**TAAAAGAGAGGCATCTTTTGAAGTAAAATTCATTTGG | Forward primer to amplify fragment F8. Used in combination with primer oDG06. *Hind*III restriction site extension |
| oDG18B | **TTTTAAGCTT**GATATAGCTCACAGTGAAACTTACTGTCAACTTGCCCC | Forward primer to amplify fragment F6B. Used in combination with primer oDG06. *Hind*III restriction site extension |

| | | |
|---|---|---|
| oDG88 | CTTGATATAGTTCACAGTGAAAG | Forward primer to amplify fragment F6C. Used in combination with primer oDG06. *Hind*III restriction site extension |
| oDG127 | TTTTAAGCTTGATATAGCTCACACTGAAAGTTAGTGTGAACTTGCCCCAT | Forward primer to amplify fragment F6D. Used in combination with primer oDG06. *Hind*III restriction site extension |
| oDG85 | GTTAGTGTGAACTTGCCCCATTTATTGTATTC | Forward primer to amplify fragment F6E. Used in combination with primer oDG86. *Hind*III restriction site extension |
| oDG86 | TTTCAGTGTGAACTATATCAAGCTTAATTGTTATCC | Reverse primer to amplify fragment F6E. Used in combination with primer oDG85. *Nhe*I restriction site extension |
| oDG146 | TTTTGGCAAGCTTGTCAGTTAATAATAAGTTTAGTA | Forward primer to amplify fragment F9. Used in combination with primer oDG147. *Hind*III restriction site extension |
| oDG147 | CGAGCAAAGCTAGCCCTAAATTTTCAATCAGTGT | Reverse primer to amplify fragment F9. Used in combination with primer oDG146. *Nhe*I restriction site extension |
| oDG11 | **TTTTAAGCT**TGTCTGTTATCTTAGTGAAAGAGTCAGTGAAAAAAATGCAG | Forward primer to amplify fragment "UP-I" region in combination with primer oDG12. *Hind*III restriction site extension. "UP-I/Down-I" fragment generate ΔUpTer deletion in pLS20 is amplify in combination with oRG17. This primer is used to generate "UP-2" (used with oDG13) in pLS20ΔUpTer deletion ("UP-2/Down-2" fragment). |
| oRG17 | **TTTTGGATCC**GGATGACAAACGATAGCAATGCCTCTGCTTGTG | Reverse primer to amplify fragments "Down-1" and "Down-2". Used in combination with primer oDG15 and oDG14B. Used in combination with oDG11 to generate fragments "UP-1/Down-1" and "UP-2/Down-2". *Bam*HI restriction site extension. |
| oDG12 | GGCAAGTTCACACTAACTTTCACTGTGAGCT | Reverse primer to amplify fragment "UP-1". Used in combination with oDG11. |
| oDG15 | **GTGAAAGTTAGTGTGAACTTGCC**TATGTTTATCAATTTGGAGCTGTCAGACTAAC | Forward primer to amplify fragment "Down-1". Used in combination with oRG17. |
| oDG13 | CCAAATTGATAAACATACATCCAAATG | Reverse primer to amplify fragment "UP-2". Used in combination with oDG11. |
| oDG14B | **GGATGTATGTTTATCAATTTGG**TTTAGGGAGGGAATAGCTTGGCTAAAGTT | Forward primer to amplify fragment "Down-2". Used in combination with oRG17. |
| Oligo 1 | GCCCTTATCCTGTTTTTAC | Competition assay for in vitro transcription. |
| Oligo 2 | GGGGCAAGTTCACACTAACTTTCACTG | Competition assay for in vitro transcription. |
| Oligo 3 | AGTCTGACAGCTCCAAATTG | Competition assay for in vitro transcription. |
| $Ter_{up28}$FW | GAACAGCTTGACAAATACACAAGAGTGTG | For amplification of $Ter_{up28}$ gBlock via PCR. |

| | | |
|---|---|---|
| $Ter_{up28}$RV | CTTTAGCCAAGCTATTCCCTCCCTAAATTTTC | For amplification of $Ter_{up28}$ gBlock via PCR. |
| $Ter_{up28}$ gBlock | GAATTCAA**GAACAGCTTGACAAATACACAAGAGTGTG**TTATAATGCAATTAGTTAATAATAAGTTTAGTAAAAACAGGATAAGGGCCATCAAAATCATGCTGCAACTTGGTTTTGATATAGCTCACAGTGAAAGTTAGTGTGAACTTGCCCCATTTATTGTATTCATACATCATTTTAGCTTTAAAAGAGAGGCATCTTTTGAAGTAAAATTCATTTGGATGTATGTTTATCAATTTGGAGCTGTCAGACTAACTGACAGCTTTTTTTGTTGTCTTTACACTGATT**GAAAATTTAGGGAGGGAATAGCTTGGCTAAAG**TTAAACGAG | gBlock to generate an in vitro transcription template containing $Ter_{up28}$. Used the gBlock as template and $Ter_{up28}$FW and $Ter_{up28}$RV as primers for amplification by PCR. Regions of primer hybridization are in bold type. |
| Nucleotides in red represents overlapping sequence. Restriction sites in lowercase. Mutations in blue. | | |
